# Functional interaction between transcription factor Sfp1 and the NuA4 complex in response to nutrient availability

**DOI:** 10.1101/2024.10.28.620578

**Authors:** Ke Xu, Stéphanie Bianco, Charles Joly Beauparlant, Valérie Côté, Lara Herrmann, Arnaud Droit, Michael Downey, Amine Nourani, Jacques Côté

## Abstract

Ribosome biogenesis is a crucial process requiring enormous transcriptional output. In budding yeast, the expression of 138 ribosomal protein (RP) genes and over 200 ribosome biogenesis (RiBi) genes is regulated by an intricate network of factors, including the nutrient-sensitive transcription activator Sfp1 and the NuA4 coactivator/acetyltransferase complex. Nutrient starvation or inhibition of TORC1 by rapamycin leads to repression of RP and RiBi genes, in part through blocking Sfp1 nuclear localization and NuA4-dependent chromatin acetylation. Here, we demonstrate that Sfp1 physically interacts with NuA4 in a TORC1-dependent manner. Our results indicate that Sfp1, along with NuA4, regulate the transcription of RiBi and RP genes via distinct mechanisms depending on promoter architectures. Sfp1 promotes histone acetylation at the promoters without affecting NuA4 recruitment. In contrast, NuA4 does impact Sfp1 binding but specifically at two classes of RP genes. Importantly, NuA4 acetylates Sfp1 at lysines 655 and 657, regulating its function. Cells expressing Sfp1 with acetyl-mimicking mutations exhibit increased expression of RiBi genes while RP genes remain stable. However, the same mutants lead to the loss of Sfp1 binding/activity at RiBi genes when cells are under non-optimal growth conditions. Mimicking constitutive acetylation of Sfp1 also limits the transcriptional burst of RP genes upon addition of glucose. Altogether, these results draw an intricate functional relationship between Sfp1 and NuA4 to control ribosome biogenesis, fine-tuning transcription output in different growth conditions.

## Introduction

Cells have evolved intricate regulatory networks to adapt their physiology to environmental changes (1). In budding yeast, the Target of Rapamycin Complex 1 (TORC1) kinase plays a central role in sensing external cues and transmitting signals to regulate cell growth and metabolism in response to nutrient availability and stress (2). Among the many physiological processes regulated by TORC1, the formation of functional ribosomes stands out as one of the most energy-intensive activities. Yeast ribosomes contain 79 different ribosomal proteins (RP) encoded by 139 RP genes. More than 200 ribosome biogenesis (RiBi) regulons are implicated in ribosome assembly to enable accurate construction and function (3). To limit unnecessary energy expenditure, the expression of RP and RiBi genes must be tightly controlled at the transcriptional level, which involves sets of activators and coactivators (4–6). Despite sharing common transcription factors (TFs) with RP genes and showing similar transcriptional responses to nutrient perturbations, RiBi gene promoters are often enriched with the so-called RRPE and PAC motifs and can exhibit distinct regulation (7).

Sfp1 is a zinc finger transcriptional activator that plays a pivotal role in orchestrating the complex expression of RP and RiBi genes in response to nutrient fluctuations (6,8). Under optimal growth conditions, TOR kinase phosphorylates Sfp1, promoting its nuclear localization, where it collaborates with various TFs and coactivators to promote RP and RiBi gene expression (9). However, nutrient starvation or treatment with rapamycin (TORC1 inhibitor) triggers the translocation of Sfp1 to the cytoplasm, resulting in RiBi and RP repression (10). Coactivators are often recruited by TFs to specific genes to modify chromatin structure, creating a permissive state for transcription. Among the coactivators implicated in RP and RiBi gene activation, histone acetyltransferase (HAT) complexes SAGA and NuA4 are recruited to the promoter region to acetylate nucleosomes and facilitate transcription (11–15). While SAGA acetylates histones H3 and H2B, NuA4 targets H4 and H2A and both complexes can be recruited through similar mechanisms (16,17). NuA4 is a 13-subunit complex that contains the only essential HAT in *Saccharomyces cerevisiae*, Esa1, required for the vast majority of H4, H2A and variant Htz1 acetylation in chromatin (18–21). We and others previously revealed the extensive binding of NuA4 at RP gene promoters, with its association seeming to be regulated by TORC1 signalling pathway (13,22–24). Esa1 positively regulates RP transcription, and its recruitment is correlated with nutrient availabilities. Together, these observations raise the question of a possible functional interaction between Sfp1 and NuA4 to regulate RP and RiBi gene expression.

To test this hypothesis and understand how these two factors may interact functionally, we undertook molecular and global approaches in budding yeast. We show that Sfp1 physically interacts with NuA4 in a TORC1-dependent manner. Depletion of Sfp1 results in reduced H3 and H4 acetylation at RiBi and RP promoters, even though NuA4 binding is not decreased at these locations. Conversely, loss of NuA4-dependent histone acetylation through Esa1 depletion affects Sfp1 recruitment at specific RP gene promoters. Interestingly, Sfp1 lysine 655 and lysine 657 are acetylated in an Esa1-dependent manner. When nutrients are available, mimicking constitutive Sfp1 acetylation leads to increased occupancy and activity at RiBi genes, whereas no significant changes are observed at RP genes. Noteworthy, cells expressing acetyl-mimicking Sfp1 are more sensitive to nutrient deficiency or TORC1 inhibition, likely reflecting misguided signaling. This implies that, in addition to histone acetylation, NuA4 regulates RP/RiBi gene expression by modulating the transcriptional function of Sfp1 through non-histone acetylation, in response to nutrient availability.

## Results

### NuA4 interacts with Sfp1 *in vivo* and *in vitro*

NuA4 and Sfp1 bind to ribosomal protein genes regulating their expression in response to stress and nutrient perturbations (13,15,24–28). However, it is still unknown whether NuA4 and Sfp1 interact physically and functionally at their mutual target genes. To answer this question, we affinity purified Sfp1-TAP fusion protein, expressed from the endogenous *SFP1* locus, and identified interacting proteins by western blotting. As a control, the NuA4 complex from an Epl1-TAP expressing strain was purified in parallel. As shown in Fig 1A, we detected the protein Tra1 in the Sfp1 purified fraction, confirming previous studies (9,26). Tra1 is an essential component of both SAGA and NuA4 histone acetyltransferase complexes. It is the recruitment interface for different transcription factors (16). To specifically ask if Sfp1 interacts with NuA4, we probed for Eaf1, a NuA4-specific subunit (29). As shown in Fig 1B, Eaf1 as well as Arp4, another NuA4 subunit, were detected in both Sfp1-TAP and Epl1-TAP purified fractions, indicating that Sfp1 interacts physically with NuA4 *in vivo*. Importantly, TORC1 inhibition with rapamycin disrupts the detected NuA4-Sfp1 interaction, likely reflecting the relocation of Sfp1 to the cytoplasm (Fig 1C). To determine if the physical interaction between Sfp1 and NuA4 can be direct, we performed a GST pulldown experiment using recombinant GST-Sfp1 and TAP-purified NuA4. NuA4 activity was assessed by a HAT assay on nucleosomes. GST-Gcn4 is used as a positive control as Gcn4-NuA4 interaction is known (30). As shown in Fig 1D, NuA4 HAT activity is pulled down by GST-Sfp1 and GST-Gcn4 but not GST alone, with parallel depletion from the supernatant. Altogether, these results indicate that Sfp1 interacts with NuA4 *in vivo* and this interaction is likely direct.

**Figure 1.**
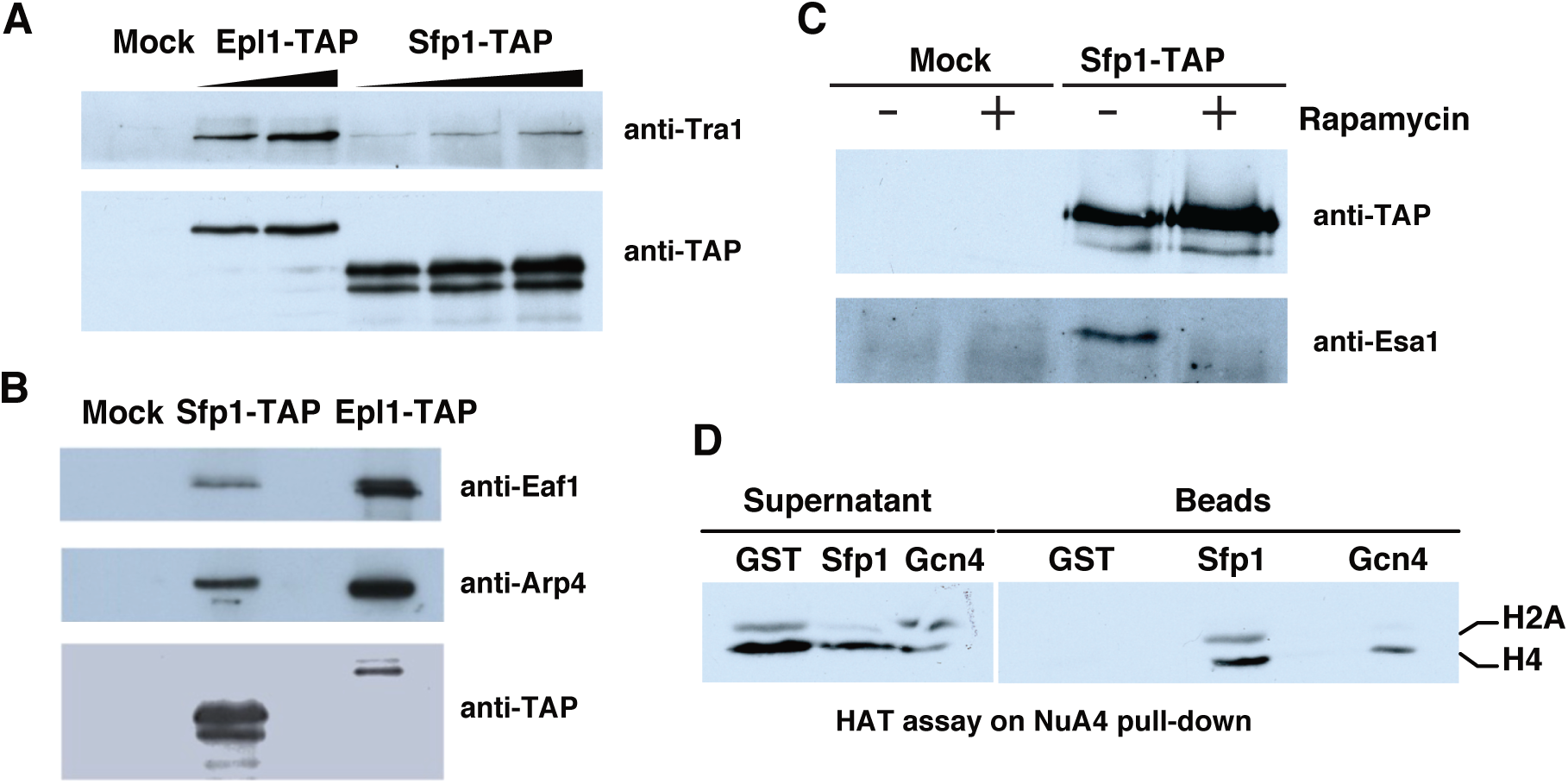
Sfp1 interacts with NuA4 in vivo and in vitro. **A)** Western blot of Tra1, a subunit of NuA4 and SAGA complexes, on different amounts of affinity-purified fractions from cells expressing Sfp1-TAP or Epl1-TAP. A mock purification from an untagged strain is shown as negative control. **B)** Western blots of Eaf1 and Arp4, selected subunits of the NuA4 complex, confirm the interaction between Sfp1 and the NuA4 complex in vivo. **C)** Sfp1 interaction with NuA4 is sensitive to rapamycin *in vivo*. Sfp1-TAP purified fractions (or mock) were obtained from cells treated or not with with rapamycin (200ng/mL) for 2 hours to inhibit the TOR kinase. Western blots of NuA4 catalytic subunit Esa1 shows loss of co-fractionating signal after treatment. **D)** TAP-purified NuA4 complex was incubated with recombinant GST, GST-Sfp1 or GST-Gcn4 proteins on glutathione beads. After washes, In vitro histone acetyltransferase (HAT) assay was performed with the beads and supernatants. HAT activity is detected on Sfp1 and Gcn4 (positive control) beads.

### Sfp1 does not affect NuA4 binding at RP and RiBi gene promoters

Since Sfp1 and NuA4 physically interact, we next determined if Sfp1 affects NuA4 recruitment at the promoters of RiBi and RP genes. Cells lacking Sfp1 exhibit reduced size and slower growth rate compared to wild-type cells (6,31,32). To avoid the negative impact of Sfp1 deletion on growth that could lead to indirect effects, we used the “anchor-away” (AA) system to rapidly deplete Sfp1 from the nucleus (33). AA-induced Sfp1 depletion results in noticeable growth defects on plates while robust growth is maintained in control conditions (Fig 2A). Sfp1 depletion from its target genes was confirmed by chromatin immunoprecipitation (ChIP) at *RPL2B* and *RPS11B* promoters (Fig S1A-B). Interestingly, the binding of NuA4 (Eaf1 subunit) does not seem affected by Sfp1 presence (Fig 2B). Even the use of *SFP1-*deleted cells confirmed that NuA4 binding at RP promoters is not significantly affected by the absence of Sfp1 (Fig S1C-D). Furthermore, even overexpression of *SFP1* from the inducible *GAL1* promoter does not affect NuA4 binding (Fig S1E-F).

**Figure 2.**
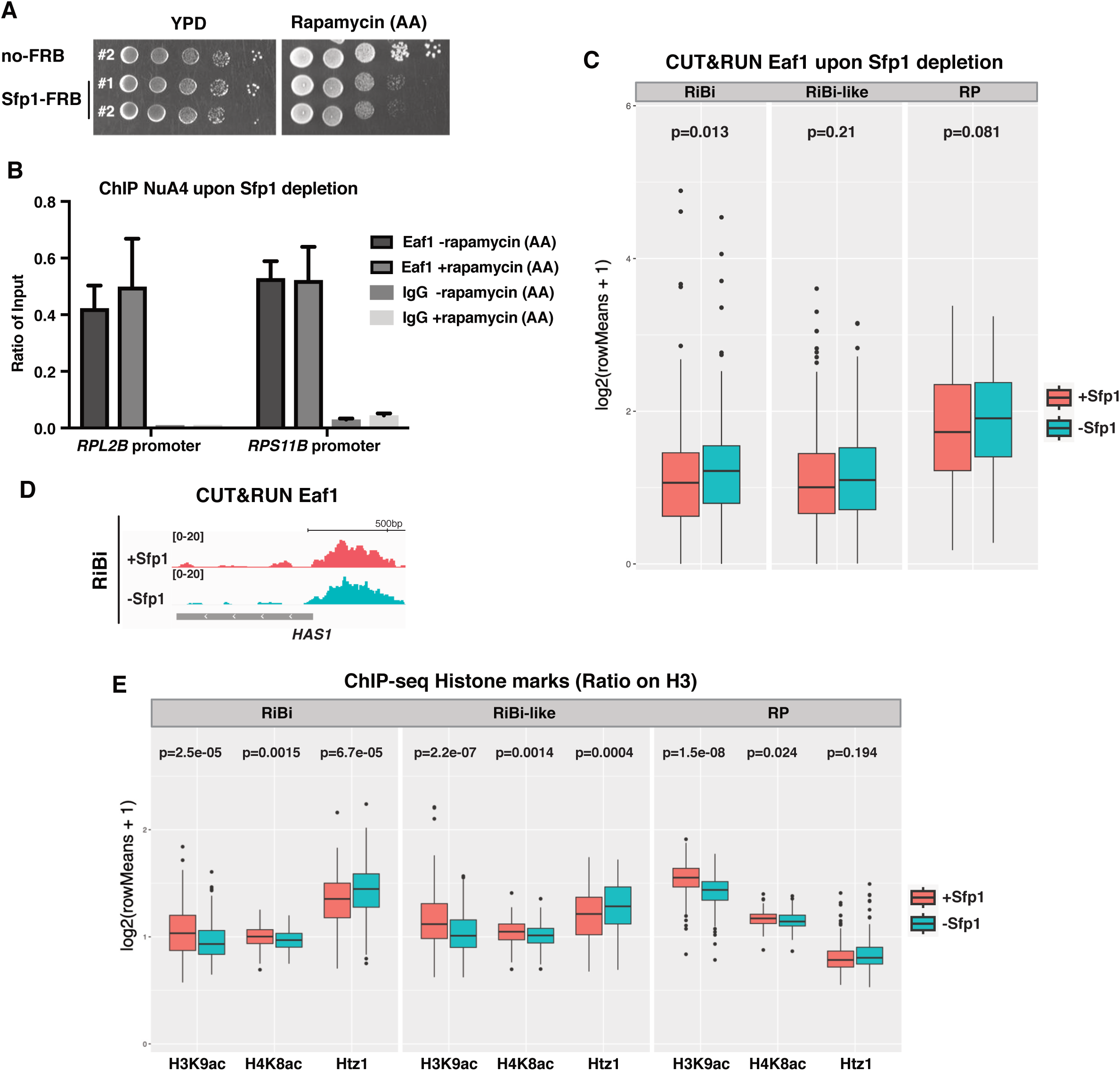
Sfp1 depletion does not decrease NuA4 recruitment on RP/RiBi gene promoters but affects chromatin acetylation. **A)** Endogenous tagging of *SFP1* with FRB was constructed in rapamycin-resistant parental strain to induce Sfp1 depletion using the anchor-away (AA) system. Rapamycin (1 µg/mL) is used to trigger AA depletion. Sfp1-depleted cells show growth defects on rapamycin agar plates whereas no-FRB control cells grow normally. **B)** NuA4 recruitment at RP gene promoters upon Sfp1 depletion was determined by ChIP-qPCR with an anti-Eaf1 antibody, showing no decrease in the absence of Sfp1. Signals are presented as ration on input (%) and Error bars are the range of two independent experiments. **C)** Anti-Eaf1 CUTnRUN samples from Sfp1-FRB cells treated or not with rapamycin were sequenced and analyzed for NuA4 localization. Box plots showing Eaf1 binding in normal conditions and upon Sfp1 depletion at the promoters of different gene clusters (Ribosomal protein (RP) genes, Ribosome Biogenesis (RiBi) genes, Ribosome Biogenesis like (RiBi-like) genes). Samples with or without Sfp1 were coloured in marron and indigo, respectively. Box plots were computed using means of bins per gene. The *p-*value was calculated by Wilcoxon testing. IgG control sequencing reads were subtracted from normalized RPM tracks. **D)** IGV tracks showing Eaf1 CUT&RUN peak at the *HAS1* gene promoter (RiBi) in normal and Sfp1-depleted cells. **E)** Box plots of ChIP-seq samples measuring histone acetylation and Htz1 occupancy changes upon Sfp1 depletion on the three gene clusters as in D. Signals were normalized to histone H3 levels measured in parallel to account for nucleosome density and are from biological replicates.

We next sought to examine Sfp1 and NuA4 localization genome-wide. Discrepancies in previous studies on Sfp1 localization stemmed from limited detection at only a subset of RP gene promoters, while both RP and RiBi genes are downregulated in *sfp1Δ* strains (6,31). Also, gene expression analysis from a Gal-induced Sfp1 expression revealed rapid induction of RiBi genes, whereas RP gene expression is increased with much slower kinetics (32). In contrast to ChIP experiments which predominantly detected Sfp1 binding at RP gene promoters, ChEC-seq using MNase-based chromatin digestion showed clear Sfp1 signals at RiBi genes with limited detection at RP gene promoters (25,34,35). This methodological discrepancy may be attributed to the higher A/T content and lower formaldehyde-crosslinking efficiency of RiBi gene promoters compared to RP genes (5,36–39). To address this issue, we used the non-crosslinking CUT&RUN approach to analyze NuA4 and Sfp1 profiles (40,41). This technique is related to the ChEC-seq method while addressing its limitations due to the target proteins being fused with the MNase enzyme. For analysis, we categorized the detectable genes into three clusters: RP genes, RiBi genes, and RiBi-like genes (other Sfp1-targeted genes that exhibit similar regulation to RiBi genes linked to growth) (25).

CUT&RUN signals for NuA4 confirm binding at both RP and RiBi gene promoters, with stronger signals observed for RP genes (Fig 2C). Consistent with our ChIP-qPCR data, Sfp1 depletion does not negatively affect NuA4 binding across all three gene clusters (Fig 2C-D). Thus, we conclude that the recruitment of the NuA4 complex is independent of its association with Sfp1.

### Sfp1 depletion attenuates acetylation of histone H3 and H4 and increases variant histone Htz1 occupancy

While the recruitment of NuA4 is not affected by Sfp1, we tested whether Sfp1 influences its function. Nucleosomes surrounding the Upstream Activating Sequences (UAS) of RP/RiBi genes are highly acetylated reflecting chromatin structure permissive for transcription (13). Histone H3 and H4 acetylation levels were measured by ChIP-seq, using total histone H3 signal as control for nucleosome occupancy, in biological duplicates. Box plots shown in Fig 2E indicate that depletion of Sfp1 induces significant reduction in both H3 and H4 acetylation at all gene clusters (also see heatmaps in Fig S2). These results suggest that the presence of Sfp1 may set the promoter architecture to facilitate histone H4 acetylation without affecting the binding of NuA4.

Htz1 is a histone H2A variant often found at promoter regions (42). It has a unique C-terminal tail that plays a crucial role in its deposition, and an extended αC helix patch that regulates chromatin compaction (43,44). Gene activation promotes Htz1 loss, whereas gene repression promotes Htz1 acquisition (42). Interestingly, Htz1 is highly enriched in RiBi genes whereas RP genes are depleted of Htz1 (42). Surprisingly, ChIP-seq data show a significant increase in Htz1 occupancy at RiBi and RiBi-like gene promoters upon Sfp1 depletion (Fig 2E, Htz1/H3). NuA4 can regulate Htz1 incorporation and is responsible for its acetylation in chromatin (21,45–47). To measure Htz1 acetylation, we immunoprecipitated proteins using an anti-acetylated lysine (anti-Kac) antibody and measured Htz1 changes by western blotting. When Sfp1 is depleted, we observe a slight decrease in Htz1 acetylation while bulk H4 acetylation remains unchanged (Fig S3A). *SFP1*-deleted cells also show a decrease in bulk Htz1K14 acetylation, as well as an apparent increase in total Htz1 (Fig S3B). Comparison of Sfp1 CUT&RUN peaks (see Fig 3 below) with published Htz1 ChIP-seq peaks (48) indicates that the majority of Sfp1 bound regions also contain Htz1 (Fig S3C). These findings are consistent with previous studies linking Htz1K14 acetylation to active genes, including Sfp1 target genes involved in ribosome biogenesis and protein synthesis (47). Our results show that Sfp1 affects NuA4-dependent acetylation of histone H4 and Htz1 as well as previously reported SAGA-mediated acetylation of histone H3 (49). Furthermore, decrease of acetylation in the absence of Sfp1 leads to stabilization of Htz1 at RiBi promoters. These observations suggest that Sfp1 indirectly modulates chromatin structure by affecting histone acetylation and the stability of Htz1-containing nucleosomes.

**Figure 3.**
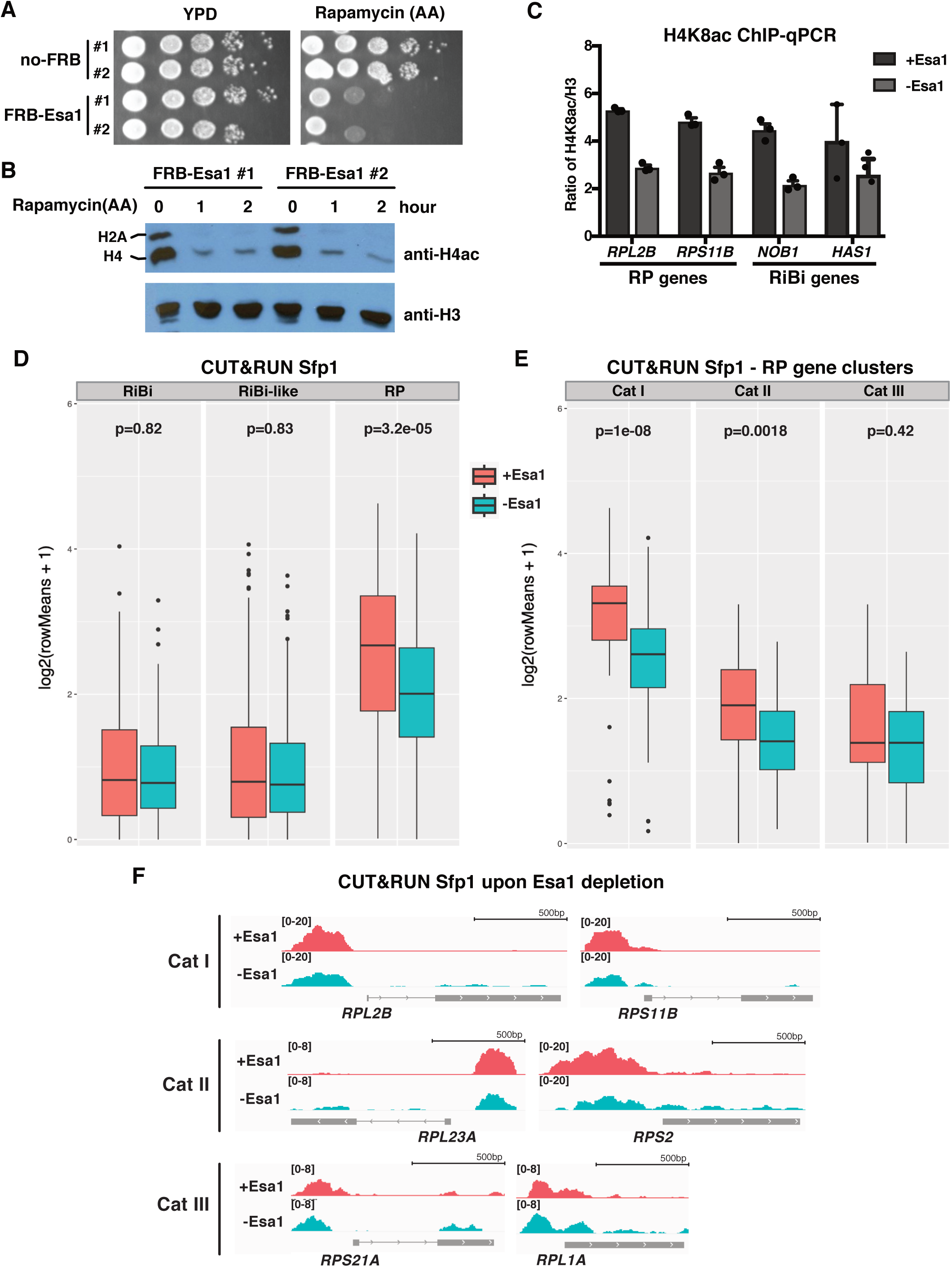
Esa1 depletion affects Sfp1 binding at specific ribosomal protein gene promoters. **A)** FRB tagging of endogenous *Esa1* was constructed in a rapamycin-resistant parental strain to induce Esa1 depletion by the anchor-away (AA) system. FRB was tagged at the N-terminus of Esa1 to avoid impeding its function. Rapamycin (1 µg/mL) is used to trigger AA depletion. Esa1-depleted cells show growth defects on rapamycin agar plates while no-FRB control cells grow normally. **B)** Rapid loss of H4 acetylation upon Esa1 depletion is shown by western blotting. The anti-H4ac antibody also recognizes H2A acetylation. Anti-H3 is used as a loading control. **C)** H4K8ac measured by ChIP-qPCR shows a decreased level at RP (*RPL2B* and *RPS11B*) and RiBi (*NOB1* and *HAS1*) gene promoters upon Esa1 depletion. Values are a ratio of IP/input of H4K8ac on H3. Error bars represent the standard error of three independent experiments. **D)** Box plots showing Sfp1 enrichment determined by CUT&RUN at the different gene clusters upon Esa1 depletion. **E)** Box plots showing Sfp1 binding at different categories of RP gene promoters upon Esa1 depletion. Box plots were computed using means of bins per gene. **F)** IGV tracks showing Sfp1 CUT&RUN peaks at different categories of RP gene promoters upon Esa1 depletion. Category I (*RPL2B* and *RPS11B*), Category II (*RPL23A* and *RPS2*), Category III (*RPS21A* and *RPL1A*). IgG controls were subtracted from normalized RPM tracks.

### NuA4 depletion leads to decreased Sfp1 binding at ribosomal protein gene promoters

Since Sfp1 does not affect the binding of NuA4 at promoters but can modulate its function, we tested if NuA4 can affect Sfp1 function at co-bound promoters. Since Esa1 is essential for viability, we relied again on the anchor-away approach to rapidly deplete NuA4 catalytic subunit Esa1 (Fig 3A). One hour of Esa1 depletion leads to reduction of H4 acetylation in bulk chromatin as well as by ChIP-qPCR at RP/RiBi genes (Fig 3B-C). We then assessed by CUT&RUN the effect of NuA4 depletion on Sfp1 presence at target genes. Upon Esa1 depletion, Sfp1 binding is reduced at the RP gene cluster (Fig 3D). In yeast, ribosomes are composed of 79 distinct RPs encoded by 138 genes. Several TFs, including Rap1, Hmo1, Ifh1, Fhl1, and Sfp1, have been found to localize at the promoter of RP genes. Distinct binding patterns of these TFs are used to categorize RP genes into three groups (3,5,50,51). Hmo1 specifically binds to Cat I promoters, while both Cat I and II promoters show detectable binding of Rap1, Ifh1, Fhl1, and Sfp1. Cat III promoters have different architectures, with only 13 RP genes in this category (39). Notably, Sfp1 has been shown to bind the gAAAATTTTc motif *in vitro* and may be present on the Cat III promoters (3,52). When Sfp1 is rapidly depleted, there is a much stronger down-regulation of Cat III genes compared to Cat I and II (25). Therefore, to more precisely assess the role of NuA4 on Sfp1 binding at RP gene promoters, we divided the RP cluster into these three categories and analyzed our Sfp1 CUTnRUN data. Box plots in Fig 3E show that Sfp1 is detectable in all three categories, although Cat III promoters had the lowest signal. Importantly, a significant decrease in Sfp1 signal is observed upon Esa1 depletion in the Cat I and II groups, whereas Sfp1 binding at Cat III genes does not seem affected. Representative IGV tracks for all three categories are shown in Fig 3F. These results clearly indicate that NuA4 is required for optimal recruitment of Sfp1 to Cat I and II RP genes.

### Sfp1 is acetylated by NuA4 at lysines 655 and 657

In addition to histones, NuA4 can acetylate non-histone substrates that are involved in a variety of cellular processes (53–56). Previous observations indicate that Sfp1 is acetylated on lysines 655 and 657 *in vivo* and this modification is Esa1/NuA4-dependent (57) (Fig 4A). To test the relative contribution of these two acetylation sites to Sfp1 function, we generated a K655/657 arginine mutant (K655/657R), which cannot be acetylated but maintains the positive charge of lysine residues, and a K655/657 glutamine mutant, which is often used as a mimic of acetylated lysine residues (K655/657Q). The R and Q mutants have the same protein level as the wild-type Sfp1 but we observed a slightly slower growth for the Q mutant in liquid media (Fig S4A-B). We first wanted to confirm that Sfp1 is acetylated *in vivo* and assess the contribution of the two lysine residues. We immunoprecipitated Sfp1 from wild-type and mutated strains and performed Western blotting using an anti-acetyllysine antibody. We also assessed Sfp1 acetylation level in a *sir2Δhst1Δhst2Δ* deacetylase mutant strain (ΔHDACs) and confirmed previous results as acetylated Sfp1 is significantly increased in that genetic background (57). As shown in Fig 4B, Sfp1 is acetylated *in vivo* and the K655/657R mutation slightly reduces the level of acetylated Sfp1 without eliminating it. This suggests that K655/657 are indeed targeted for acetylation but are not the only sites on Sfp1. It also indicates that these sites show dynamic acetylation/deacetylation cycles involving sirtuins deacetylases (Fig 4B).

**Figure 4.**
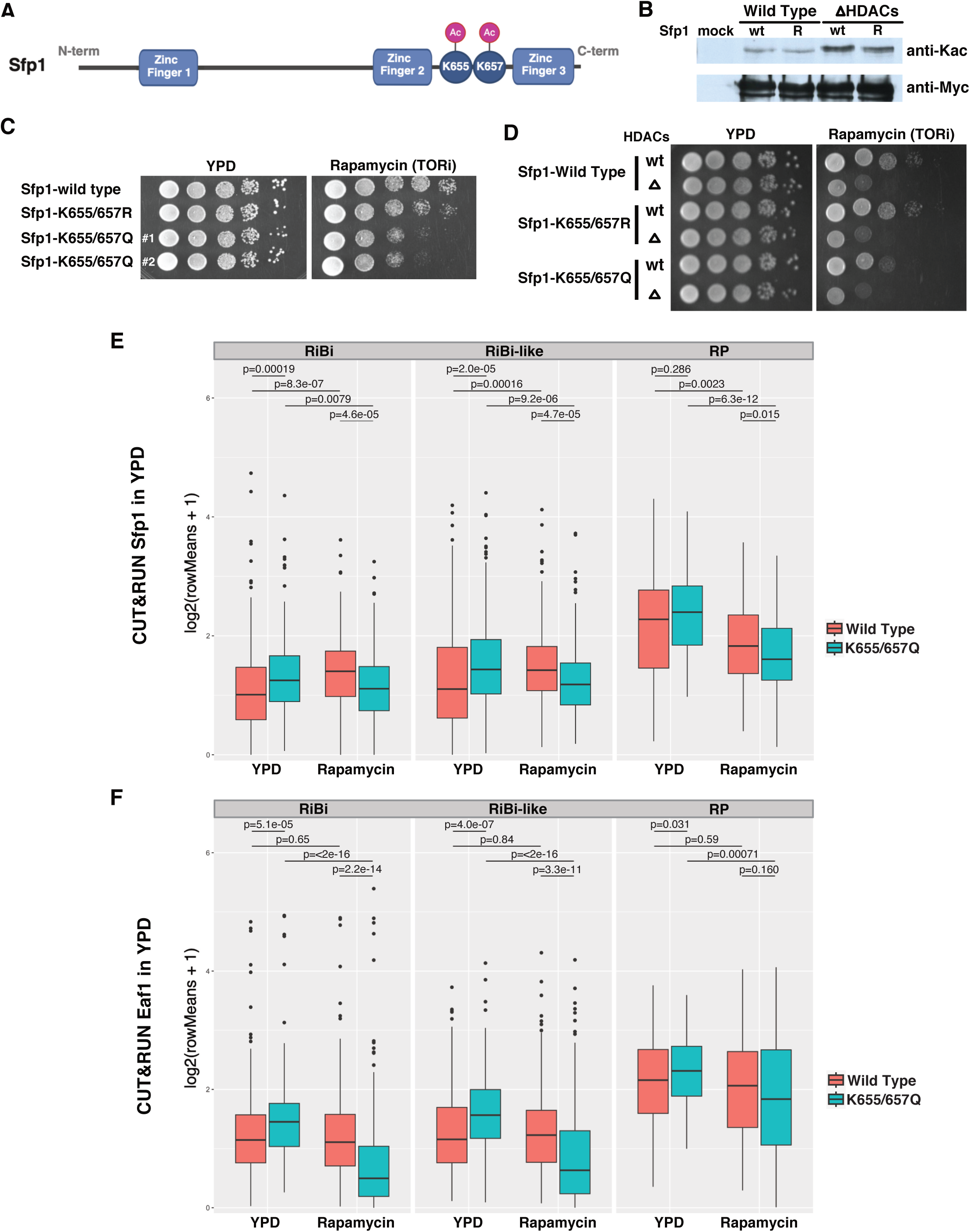
Esa1-dependent acetylation of Sfp1 on lysines 655 and 657 affects its binding at target genes in normal and TOR-inhibited conditions. **A)** Diagram showing the locations of the lysine residues acetylated by Esa1 and the predicted Zinc-finger domains of Sfp1. **B)** Sfp1 acetylation is regulated by sirtuin histone deacetylases. Myc-tagged Sfp1 was Immunoprecipitated from wild-type or *sir2*Δ *hst1*Δ *hst2*Δ (ΔHDACs) triple mutant cell extracts and analyzed by western blotting with an antibody directed towards acetylated lysine or the myc epitope. The acetylation status was compared between wild-type Sfp1 and Sfp1 with lysines 655 and 657 mutated to arginine (WT vs R). **C)** Spot assay testing growth rates of wild-type cells and Sfp1 mutants in YPD and Tor-inhibited conditions. Ten-fold serial dilutions of strains expressing WT or mutant Sfp1 on complete media alone or containing rapamycin (200ng/mL). **D)** ΔHDACs cells are hypersensitive to rapamycin. Sfp1 K655/657 mutants were introduced in *sir2*Δ *hst1*Δ *hst2*Δ (ΔHDACs) triple mutated strain and spot assays were performed as in C. **E-F)** Box plots showing Sfp1 and Eaf1 CUT&RUN signals at different gene clusters in the wild-type and K655/657Q mutant strains, in normal rich media (YPD) and TOR-inhibited (200ng/mL rapamycin) conditions. Signals obtained in parallel with IgG control were subtracted from normalized RPM tracks. Wild-type Sfp1 is coloured in marron, K655/657Q mutant coloured in indigo. Box plots and associated Wilcoxon tests were computed using means of bins per gene.

In optimal growth conditions, Sfp1 is localized in the nucleus and moves out to the cytoplasm when yeast cells undergo stress or nutritional deprivation (10). As mentioned above, the TORC1 Complex promotes Sfp1 nuclear localization (6) and its inhibition by rapamycin strongly affects Sfp1 localization and function. Interestingly, the Sfp1 K655/K657Q acetyl-mimic mutant cells have increased sensitivity to rapamycin compared to wild-type and K655/657R mutant cells (Fig 4C). In parallel, the ΔHDACs strain is highly sensitive to TORC1 inhibition but the K655/657R mutant cannot rescue this phenotype by itself (Fig 4D). These results suggest that mutations in Sfp1 mimicking constitutive acetylation makes the cells more sensitive to inhibition of TOR signaling, perhaps acting like a misleading signal of high nutrient conditions. To dissect the effects of NuA4-dependent Sfp1 acetylation on Sfp1 binding to target genes, we performed Sfp1 CUT&RUN sequencing under normal and TORC1-inhibited conditions.

Intriguingly, the binding of the acetyl-mimic Sfp1 is significantly increased at RiBi and RiBi-like genes when growth conditions are favourable, while no significant change is observed at RP gene promoters (Fig 4E, S4C). In contrast, the opposite trend occurs upon rapamycin treatment, as the acetyl-mimic Sfp1 shows significant decreased binding (Fig 4E). This is consistent with the slower growth rate in the presence of rapamycin, the acetyl-mimic mutant being more sensitive to TORC1 inhibition than wild-type Sfp1. Importantly, NuA4 recruitment is also increased at RiBi and RiBi-like gene promoters in rich media in the presence of Sfp1-K655/657Q as well as also decreased after rapamycin treatment in the same background, functionally linking again Sfp1 acetylation and NuA4 (Fig 4F). Altogether, these results suggest that constitutive acetylation of Sfp1 is an aberrant signal misleading the cell in different growth conditions.

### Acetylation of Sfp1 affects its activity as transcriptional activator

Since the binding of Sfp1 to target genes can be affected by its NuA4-dependent acetylation, we analyzed its effect on transcription. RNAs were extracted from wild-type Sfp1 and the two mutant strains grown in rich media and differentially expressed genes were measured by RT-qPCR. The K655/657Q strain shows more than two-fold increase in the expression of RiBi genes, while RP gene expression remains similar to that of the wild-type strain (Fig 5A). To get the genome-wide picture, we then performed RNA sequencing (Fig S5). As shown in Fig 5B-D, genes belonging to RiBi and RiBi-like clusters are confirmed as upregulated in the Sfp1 acetyl-mimic mutant. Altogether, these results demonstrate that NuA4-dependent acetylation of Sfp1 has a significant impact on its binding and function regulating specific classes of genes. This is also reminiscent of previous observations showing that overexpression of Sfp1 upregulates a large number of genes involved in ribosome biogenesis, leading to a slow-growth phenotype (8). Similarly, the same set of genes related to metabolic pathways is down-regulated in K655/657Q mutant and Sfp1-overexpressing cells (Fig S6) (25).

**Figure 5.**
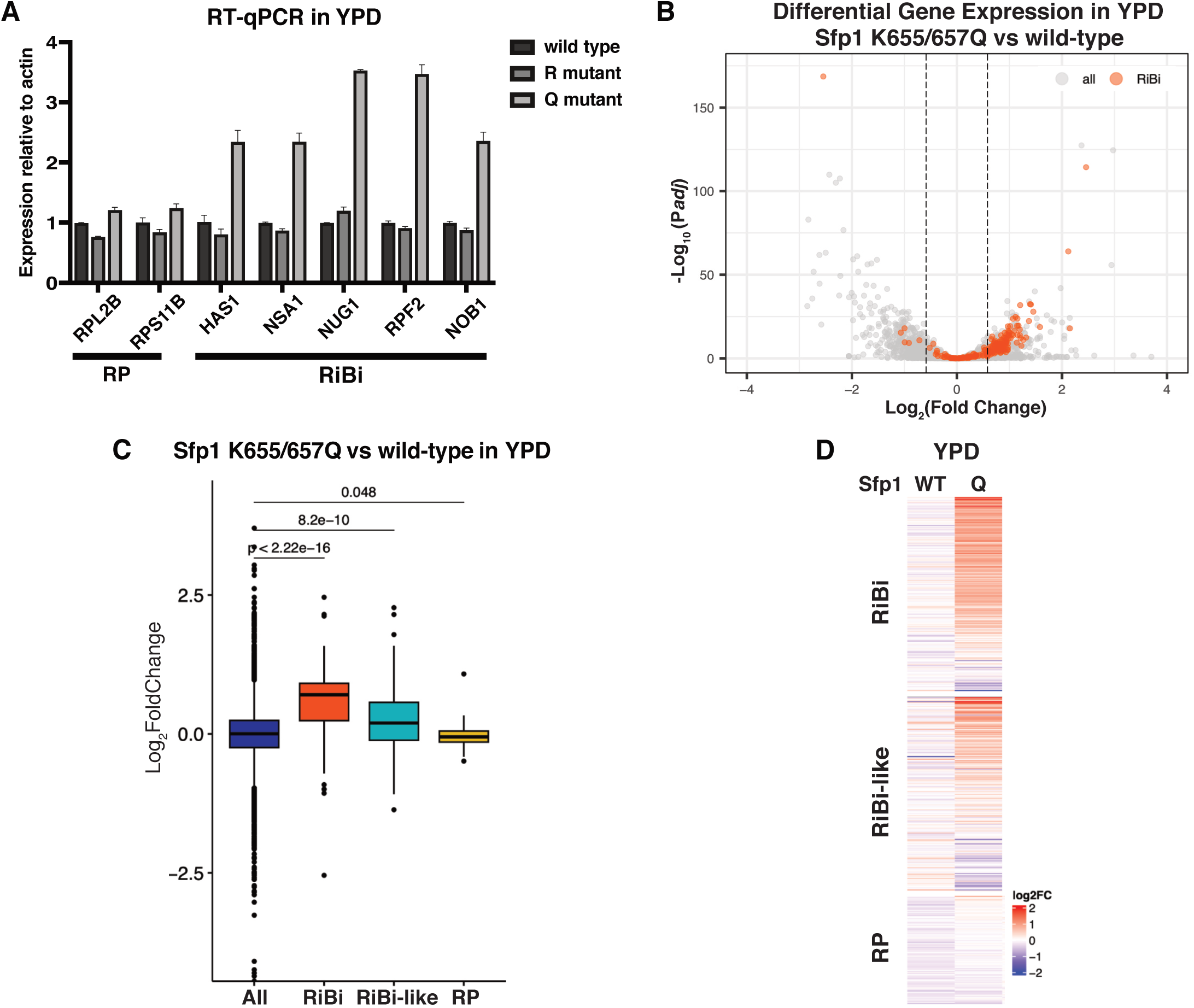
Acetylation-mimic Sfp1 K655/657Q mutant upregulates RiBi gene expression under normal growth conditions. **A)** RNAs were extracted from log phase wild-type Sfp1, K655/657R, and K655/657Q strains growing in YPD. mRNA level of indicated RiBi genes (*NOB1*, *HAS1*, *NUG1*, *RPF2* and *NSA1*) and RP genes (*RPL2B* and *RPS11B*) were quantified, relative to levels of *ACT1* mRNA, by RT-qPCR. Error bars represent the range of two independent experiments. **B)** Volcano plot of differential gene expression analysis by RNA-seq of the Sfp1-K655/657Q mutant strain compared with wild-type. RiBi genes are highlighted in orange. Genes were considered significantly dysregulated for absolute fold change ratio >=1.5 and adjusted p-value <0.05. **C)** Box plots showing gene expression changes of different clusters (RiBi, RiBi-like, and RP genes) in K655/657Q mutant compared with wild-type cells growing in YPD. **D)** Heatmap showing differential gene expression of the wild-type strain compared with the acetyl-mimic Sfp1 strain.

### Carbon source shift and glucose pulse experiments reveal different Sfp1 regulatory mechanisms toward RiBi and RP gene clusters

Yeast cells deleted for *SFP1* are characterized by carbon source-dependent slow growth and small sizes (31,32). While glucose is the preferred carbon source, yeast can also ferment raffinose as a non-permissive carbon source, but at a lower growth rate. We repeated RNA extraction, RT-qPCR and RNA-seq with the wild-type and mutant strains but this time in raffinose-containing media (Fig 6, S5). When growing in the media where raffinose is the sole carbon source, RP genes are downregulated in all tested strains. However, RiBi genes seem to some extent upregulated (Fig 6A). Strikingly, with this carbon source, an opposite effect is seen with the Sfp1 acetyl-mimic mutant, with RiBi and RiBi-like genes being down-regulated (Fig 6A-D). These results are certainly consistent with the Sfp1 CUT&RUN data showing that binding of the acetyl-mimic Sfp1 is more sensitive to stress/nutrient starvation (Fig 4E).

**Figure 6.**
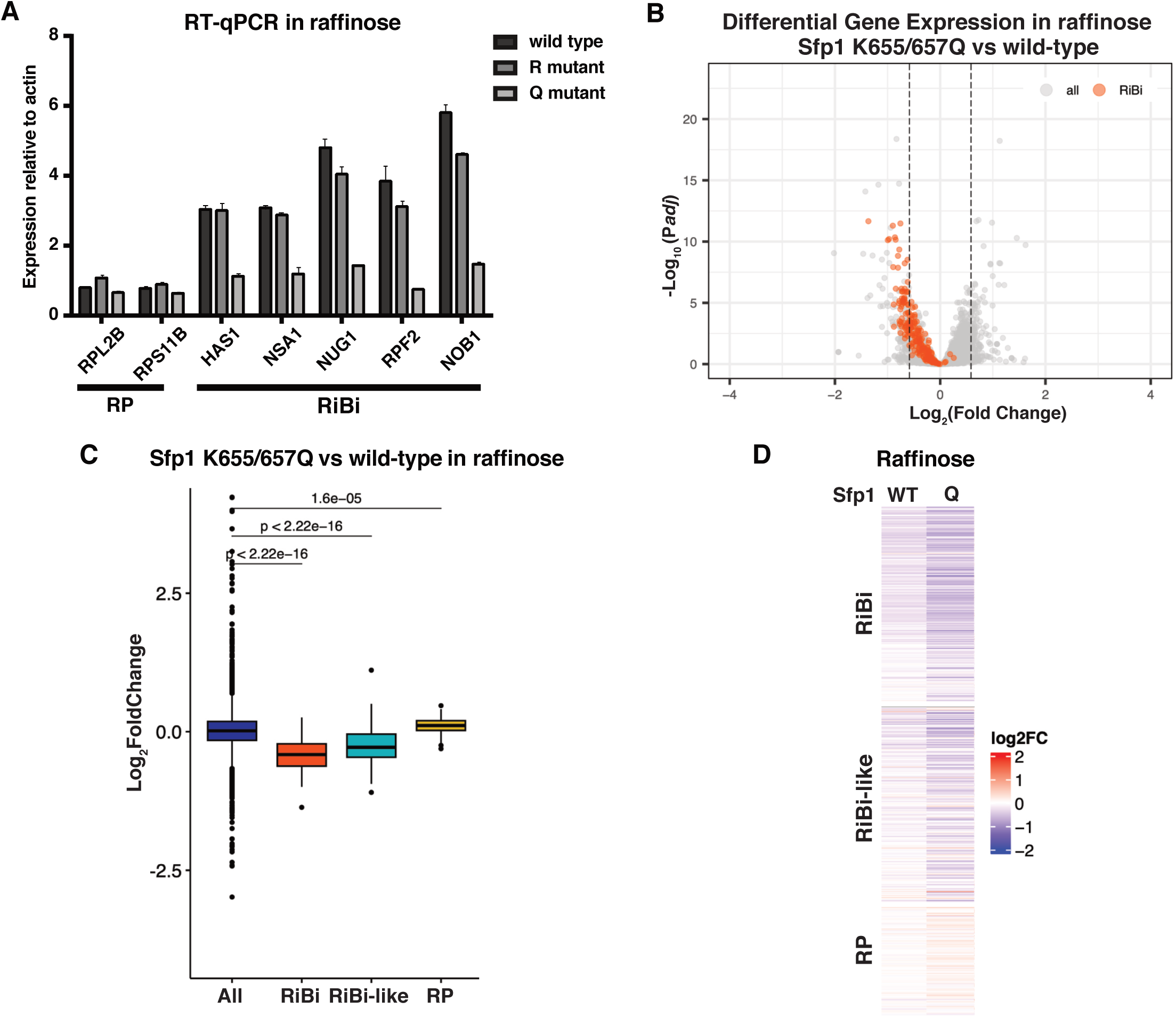
K655/657Q mutations in Sfp1 create sensitivity to carbon shift from glucose to raffinose. **A)** Cells expressing indicated Sfp1 and its mutations were grown in YPD to log phase before transferred into media that contains raffinose as single carbon source. mRNA level of indicated RiBi genes (*NOB1*, *HAS1*, *NUG1*, *RPF2* and *NSA1*) and RP genes (*RPL2B* and *RPS11B*) were quantified, relative to levels of *ACT1* mRNA, by RT-qPCR. Error bars represent the range of two independent experiments. **B)** Volcano plot of differential gene expression analysis by RNA-seq of the Sfp1-K655/657Q mutant strain compared with the wild-type strain, grown in raffinose. RiBi genes are highlighted in orange. Genes were considered significantly dysregulated for absolute fold change ratio >=1.5 and adjusted p-value <0.05. **C)** Box plots showing gene expression changes between different clusters (RiBi, RiBi-like, and RP genes) in K655/657Q mutant compared with wild-type cells grown in raffinose. **D)** Heatmap showing differential gene expression in raffinose of wild-type strain compared with acetyl-mimic Sfp1 strain.

To better understand the different effects of carbon sources and eliminate growth rate influences, we conducted a carbon source shift assay, adding glucose back after incubation in raffinose. This assay allowed us to study how cells detect changes in nutrient availability and adjust their metabolism. Fig 7A illustrates the procedure of the assay. Both K655/657 mutants (R and Q) and reference strains were grown in rich YPD media (2% glucose) until the early exponential phase. Cells were then washed to remove glucose and incubated in YPR (2% raffinose) for one generation. Glucose was then reintroduced to the media and gene expression was monitored at different time points.

**Figure 7.**
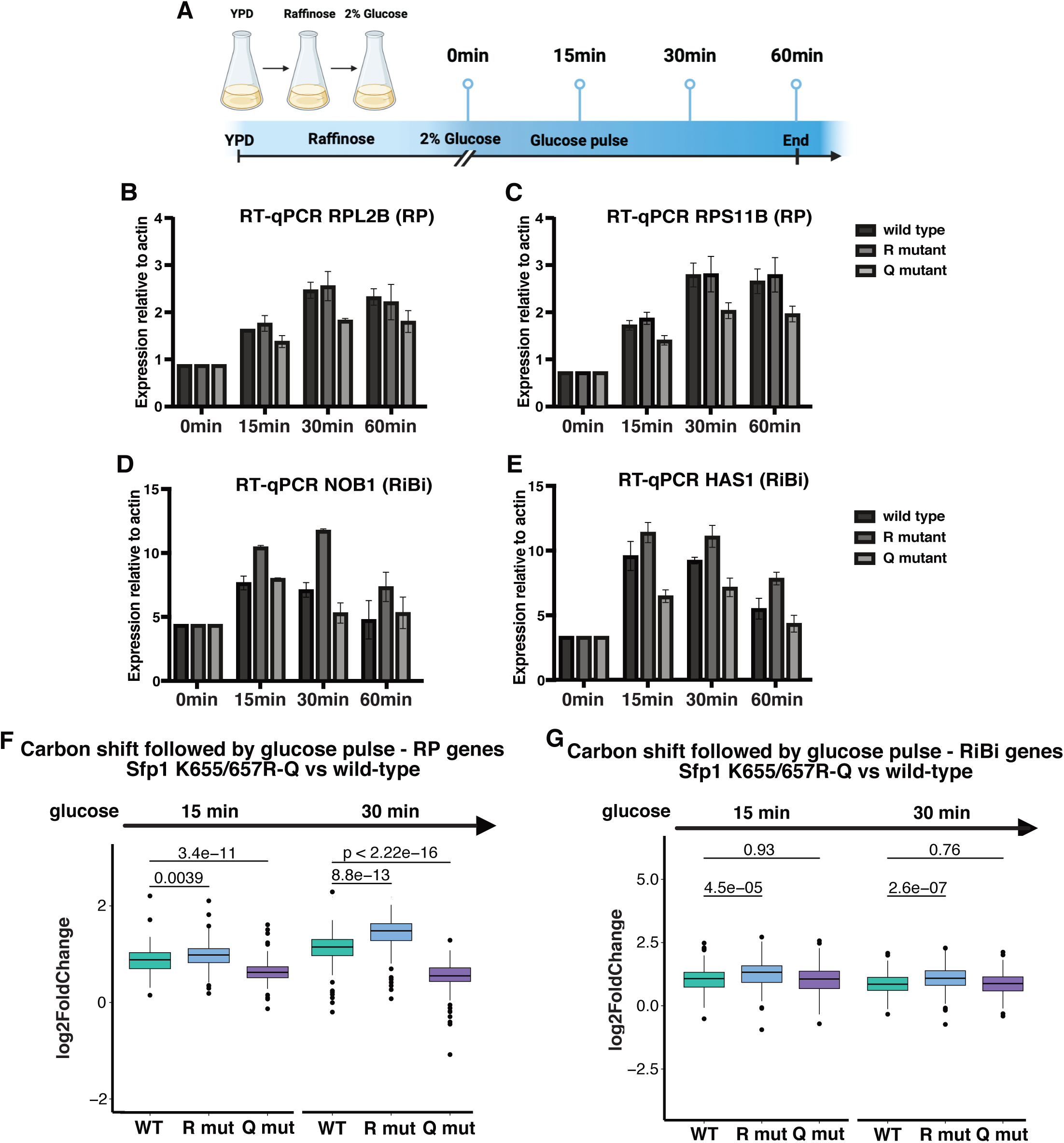
NuA4-dependent acetylation of Sfp1 regulates transcription of RP genes differently than RiBi genes upon a glucose pulse after the shift in carbon source. **A)** Flow diagram of the carbon shift assay. Wild-type and Sfp1 mutants were grown in raffinose prior to the addition of glucose. Cells were collected at different time points for RNA quantifications. **B-E)** RNAs were extracted from log phase wild-type Sfp1, K655/657R, and K655/657Q strains growing in the indicated conditions. Gene expression levels of selected RP genes (*RPL2B* and *RPS11B*) and RiBi genes (*NOB1* and *HAS1*) were quantified, relative to levels of *ACT1* mRNA, by RT-qPCR. Values were normalized to time 0 in each strain. Error bars represent the range of two independent experiments. **F-G)** Box plots showing gene expression changes determined by RNA-seq upon the glucose pulse comparing the wild-type and Sfp1 mutant strains. Glucose was added back into the media for the indicated times and values were normalized to the 0 timepoint. RiBi and RP gene clusters are shown separately.

As expected, RT-qPCR data show that the expression of both RP and RiBi genes increases immediately upon the addition of glucose (Fig 7B-E, S7A). The Sfp1 K655/657R mutant and wild-type strains show similar expression levels of tested RP genes during the glucose pulse period, while RiBi genes seem slightly more expressed in the K655/657R mutant. However, global analysis by RNA-seq reveals that the K655/657R mutant, while confirming more expression of RiBi genes, also shows significant more expression of RP genes compared to wild-type cells (Fig 7F-G). These results indicate that, although the K655/657R mutant does not show growth phenotype under normal conditions, Sfp1 function is still affected under poor conditions when cells are suddenly pushed to their potential by rapid transcriptional reprogramming upon addition of optimal carbon source. In parallel, despite the RiBi gene expression differences compared to wild-type cells, the acetyl-mimic Sfp1 mutant is able to drive RiBi gene transcription to a level similar to wild-type cells upon the glucose pulse (Fig 7G). Interestingly, RP gene upregulation with the addition of glucose shows a clearly different regulatory mechanism since the K655/657Q mutant does not show capacity to express RP genes to the same level as the wild-type cells during the glucose pulse (Fig 7F, S7B).

## Discussion

In the present work, we demonstrate that the Sfp1 transcription activator interacts with the NuA4 coactivator complex to coregulate ribosomal protein and ribosome biogenesis gene expression. The presence of Sfp1 at the gene promoters is important for the NuA4-mediated local histone H4 acetylation. Despite NuA4 binding to promoters not being affected by Sfp1, its function as acetyltransferase seems attenuated with Sfp1 depletion. We also revealed that histone variant Htz1 occupancy increases whereas its acetylation levels may decrease upon Sfp1 depletion. NuA4-dependent Htz1 acetylation near gene promoters was reported to be linked to transcription activation since it favors nucleosome disassembly (42,47). These results indicate that Sfp1 may facilitate gene transcription via NuA4-dependent Htz1 acetylation. Our previous work showed that NuA4-mediated H4 acetylation can increase Htz1 incorporation which could also be at play in NuA4 and Sfp1 functional interaction (46). In addition, NuA4 has a clear positive impact on Sfp1 binding at RP genes in optimal growth conditions, indicating bi-directional functional cooperation (Fig 8A).

**Figure 8.**
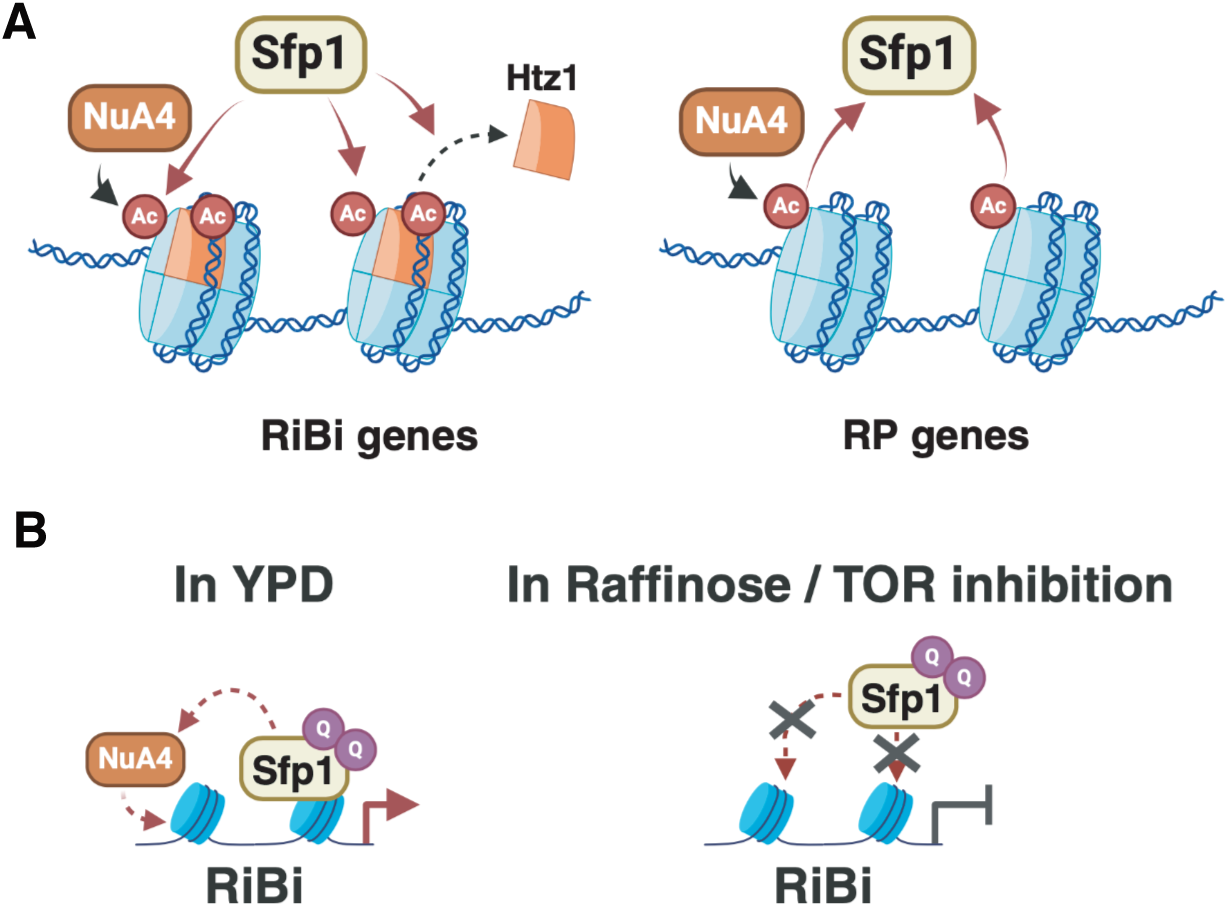
Interplay of NuA4 and Sfp1 at RP and RiBi genes. **A)** In rich media Sfp1 is not involved in NuA4 recruitment but affects its acetyltransferase function at RiBi/RP gene promoters. In parallel, NuA4 stimulates the binding of Sfp1 at RP gene promoters. **B)** Mimicking constitutive acetylation of Sfp1 by NuA4 stimulates its binding to RiBi genes in rich media but is detrimental in stress/poor carbon source conditions.

The physical interaction described here between NuA4 and Sfp1 is also reminiscent of the protein JAZF1, the homolog of Sfp1 in mammals. JAZF1 is strongly linked to cellular metabolism, being implicated in glucose and lipid homeostasis, insulin signaling, and metabolic disorders such as diabetes (58). Strikingly, JAZF1 has recently been identified as a stoichiometric subunit of the human NuA4/TIP60 complex (59). Furthermore, the depletion of JAZF1 can lead to reduced H2A.Z acetylation at NuA4/TIP60 regulatory sites (60). Thus, the physical and functional interaction between NuA4 and Sfp1 seems conserved throughout eukaryotes and play a major role in regulating cell homeostasis.

The fact that Sfp1 is directly acetylated by NuA4 is certainly part of signaling events regulating its function upon changes in growth conditions. Cells expressing acetyl-mimic mutant Sfp1 exhibit abnormal transcription profile that includes augmented RiBi gene expression under normal growth conditions and reduced RiBi gene expression under stress/poor carbon source, both linked to level of Sfp1 binding to the promoters (Fig 8B). The high availability of acetyl-CoA in rich media may be part of the process leading to Sfp1 acetylation by NuA4, signaling optimal conditions for growth, hence ribosome biogenesis/translation. But this possible link with acetyl-CoA does not fit with how the acetyl-mimic mutant misleads the cells in poor conditions, in this case seemingly overcompensating by showing reduced expression of RiBi genes. One model to reconcile those results could argue that cycles of Sfp1 acetylation and deactylation serve as a rheostat to ensure proper transactivator activity, not too high or too low. Due to the complexity of RP gene regulation under steady state conditions, only an impact of the Q mutant on RiBi genes can be seen, but contributions of both K-R and K-Q mutants are seen during nutrient shifts. Importantly, in these shifting conditions, the mutants show opposite effects on RP genes, as one might predict (Fig 7F).

The glucose pulse experiment allowed us to explore different transcriptional mechanisms between RiBi genes and RP genes mediated by Sfp1, before other mechanisms of regulation kick in at steady state levels. As mentioned before, RiBi gene promoters have two distinct motifs indicating that RiBi gene regulations are partially different from RP genes. Also, transcription factor Ifh1, which binds RP gene promoters, affects Sfp1 binding only at RP genes (25) and is itself regulated by acetylation, although through Gcn5/SAGA instead of Esa1/NuA4 (12). Interestingly, Ifh1 acetylation titrates its activator function after being bound the RP promoters and it is tempting to speculate a related fine-tuning function of Sfp1 acetylation by NuA4 at RiBi genes. Constitutive Sfp1 acetylation may lead to aberrant cell response to nutrient perturbations through specific regulatory mechanisms towards different targeted gene clusters. A deeper mechanistic understanding will require precise measurement of Sfp1 acetylation in different conditions as well as characterizing other lysine residues being targeted. In addition, coordinated analysis of functional interplays of Sfp1 with both NuA4 and SAGA acetyltransferase complexes, in response to growth condition changes and stress, will shed more light on an undoubtedly very complicated regulatory network.

## Acknowledgments

We are grateful to Steven Henikoff for providing Protein A-MNase used in CUT&RUN experiments. This work was supported by grants from the Canadian Institutes of Health

Research (CIHR) to J.C. (FDN-143314, PJT-183708). K.X. was supported by a scholarship from the Chinese Academy of Science. J.C. held a Canada Research Chair (Tier 1) in Chromatin Biology and Molecular Epigenetics.

## EXPERIMENTAL PROCEDURES

### Yeast strains and spot assay

Yeast strains used in this study are listed in S1 Table and were constructed based on standard PCR-based transformation protocol. All strains generated were verified using PCR and sequencing. Unless indicated, Yeast cells were grown in YPD (1% yeast extract, 2% peptone, 2% dextrose) at 30°C. The anchor-away parental strain used for transformation was previously mutated (*tor1-1* mutation and *FPR1* deletion) to make the cells resistant to rapamycin, while Rpl13a was tagged with FKBP12, as described (33). For Esa1 N-terminal tagging, the FRB tag was inserted after the start codon as described (61). Cells were treated with rapamycin at 1 μg/mL for anchor-away-induced depletion. For spot assays, single colonies from fresh plates were grown overnight in YPD at 30°C. Cell cultures were diluted in fresh YPD to OD_600_ 0.3 and continued to grow until OD_600_ 0.7-0.8. The 10-fold serial dilutions of the yeast cultures were made for each strain and 4 μL of each dilution was spotted on plates. Colonies were incubated at 30°C for 2-4 days, then the growth sensitivities were compared between plates.

### GST purification and HAT assay

Recombinant Sfp1 was purified from BL-21 bacteria transformed with pGEX-4T-3-GST-*SFP1* and induced with 500 nM IPTG overnight at 16°C. Bacterial pellets were lysed with lysozyme and sonicated. After clarification by centrifugation, soluble fractions were incubated with Glutathione-Sepharose beads (GE Healthcare) for 3 hours at 4°C. Beads were washed and ready to use. GST-immobilized Sfp1 proteins were incubated with TAP-purified NuA4 complex for 4 hours at 4°C. After incubation, the supernatant was collected, and the beads were washed. Histone acetyltransferase (HAT) assay was performed as described (62,63). 500ng of short oligonucleosomes purified from MNase-digested yeast chromatin was incubated with the GST beads or the supernatant mentioned before at 30°C for 30 minutes. 0.125 μCi of [3H] acetyl coenzyme A ([3H] acetyl-CoA) was added in HAT buffer (50 mM Tris-HCl [pH 8], 50 mM KCl plus NaCl, 0.1 mM EDTA, 5% glycerol, 1 mM DTT, 1 mM PMSF and 20 mM sodium butyrate). The reactions were stopped in 1X Laemmli buffer, boiled and loaded on a 10% SDS-PAGE. Gel was treated with En3hance (Perkin Elmer), dried and exposed on film.

### Chromatin immunoprecipitation (ChIP)

200 mL cell cultures were collected at OD_600_ 0.7-0.8. Cells were crosslinked with 1% formaldehyde for 20min and quenched by adding 125 mM glycine for 5min. Sonication was performed to obtain 200-500bp chromatin fragments. 100 ug of chromatin was incubated with indicated antibodies for each IP. Magnetic beads were added, and the mix was incubated for 3-4 hours at 4°C. The chromatin was eluted from beads after wash. The eluate was incubated overnight at 65 °C to reverse the crosslinks. Phenol chloroform extraction was performed and 1uL of each DNA sample was used for qPCR. ChIP-qPCR data are presented as % of IP/Input (or ratio of IP/Input when correcting for nucleosome occupancy, i.e. H4K8ac/H3) and are from at least two independent yeast cultures in each experiment. Lists of primers standardized on a LightCycler qPCR apparatus are available upon request. For ChIP-sequencing, libraries were prepared as described (64). Samples were sequenced 100-bp paired-end for around 10M reads per sample. Sequencing was performed at IRCM (Montreal Clinical Research Institute). Antibodies used were against H4K8ac (Abcam, ab45166), H3K9ac (Upstate, #07-352), Htz1.K14ac (Upstate, #07-719), H3 (Abcam, ab1791), Htz1 (Millipore, #07-718), Eaf1(homemade), Myc (Sigma, M5546).

### Acetyl-lysine Immunoprecipitation

Immunoprecipitation was performed as described previously (65). Cells were grown in YPD till OD600 around 0.5 followed by rapamycin (1 μg/mL final) or vehicle treatment for 2 hours. Cells were collected and lysed in lysis buffer (10 mM Tris-HCL pH 8.0, 150 mM NaCl, 10% glycerol, 0.1% NP-40, 2 μg/mL leupeptin, 2 μg/mL pepstatin A, 5 μg/mL aprotinin, 1 mM PMSF, 10 mM *β*-glycerophosphate, 1mM Sodium Butyrate, 0.5 mM NaF, and 1 mM DTT). 6 μg of WCE was incubated with anti-acetyl (ImmuneChem ICP0380) overnight at 4°C. Protein G magnetic beads (Invitrogen 1004D) were added and incubated for 4 additional hours. Beads were washed and 1X Laemmli buffer was added. The samples were then loaded on SDS-PAGE.

### RT-qPCR and RNA-sequencing

Total RNA was extracted following the hot-phenol protocol. In brief, cell pellets were resuspended in 400 μL TES solution (10 mM Tris HCl, 10 mM EDTA, and 0.5% SDS). 400 μL acid phenol was added and then vortexed. The cell suspensions were incubated at 65 ° C for 30 minutes with occasional, brief vortexing. Top speed centrifuge was performed to separate the aqueous phase. The aqueous phase was transferred and 400 μL chloroform was added followed by another round of vortex and centrifuge. RNAs were precipitated with sodium acetate and 100% ethanol. Initial precipitated RNAs were washed with ice-cold 70% ethanol. RNA pellets were resuspended in 50 μL RNase-free H_2_O. Reverse transcription was performed following the 2X SYBR Green PCR Master Mix kits protocol. qPCR was performed using indicated primers to test specific gene expression and normalized to ACT1 expression level. For RNA-Seq, the NEBNext ultra II RNA library prep kit was used for library preparation. The sequencing run was performed on an Illumina NovaSeq 6000 system. Reads were trimmed using fastp v0.23.2 (66). Quality check was performed on raw and trimmed data to ensure the quality of the reads using FastQC v0.11.9 and MultiQC v1.12 (67). The quantification was performed with Kallisto v0.48.0 (68) against the Saccharomyces cerevisiae transcriptome (R64-1-1 downloaded from Ensembl release 109). The principal component analysis was completed with the FactoMineR v2.7 R package (69) and the graphical representations were produced with the ggplot2 v3.4.2 package (70). The sequencing data of K655/657R-2 at the 30-minute time point has been removed according to the PCA as an outlier sample (See Fig S5). Differential expression analysis was performed using the DESeq2 v1.38.3 package (71). Genes were considered significantly dysregulated for absolute fold change ratio >=1.5 and adjusted p-value <0.05. The volcano graphical representations were produced with the EnhancedVolcano v1.18.0 package. GO enrichment analysis was carried out using clusterProfiler v4.8.3 with an adjusted p-value cutoff of 0.05 and the function “simplify” to reduce the redundancy of GO terms enriched with the default parameters (72). Box plots and associated Wilcoxon tests were generated using the ggboxplot and stat_compare_means functions, respectively, with the default settings of the ggpubr v0.5.0 package (https://CRAN.R-project.org/package=ggpubr). Heat maps were generated using ComplexHeatmap v2.10.0 package (73). All R analysis were done in R v4.2.2.

### CUT&RUN Sequencing

The experiment was performed as described (41). Yeast nuclei were freshly prepared according to published procedures (74) and bound to concanavalin A beads. The beads were then resuspended in a buffer containing digitonin, and antibodies (Eaf1, Myc and IgG) were added to incubate overnight at 4°C. Afterward, the beads were washed, and pA-MNase (provided by Steven Henikoff) was added. One-hour secondary antibody incubation was performed for samples using anti-Myc antibody. CaCl2 was added to activate MNase digestion for 5 to 15 minutes. The reaction was quenched by the addition of stop buffer containing EDTA and EGTA. DNA was purified with NEB Monarch PCR and DNA purification kit following the protocol. DNA was quantified by Qubit HS DNA kit. The library was prepared following the manufacturer’s instructions in NEBNext Ultra II DNA kit for low-input ChIP. 100-bp pair-end sequencing was performed with the Illumina NovaSeq 6000 system.

### CUTnRUN/ChIP-seq analysis

Raw reads were trimmed using fastp v0.20.1 (66). Trimmed reads were aligned on the Saccharomyces cerevisiae genome (R64-1-1 - sacCer3) using bwa mem v0.7.17 (75) and samtools v1.13 (76). The macs2 v2.2.1 software (77) was used to perform the peak calling and regions were annotated using the ChIPseeker package v1.30.3 (72) in R v4.1.2. Raw signal tracks and normalized tracks (in reads per million, RPM) were produced from mapped reads using deepTool’s bamCoverage tool v3.3.0 (78) and bedtools genomecov tool v2.17.0 (79), respectively. Tracks were converted to the bigwig format using bedGraphToBigWig v2.8 (80) and signal tracks were visualized with IGV (81). For transcription factors, IgG negative control tracks were subtracted from normalized RPM tracks using deepTools ‘bigwigCompare’ (78). For histone marks, signal tracks were normalized to histone H3 levels. Reads mapping were then divided into promoter groups of genes (RIBI, RIBI-like, RP genes and subcategories) for the generation of heat maps using ComplexHeatmap v2.10.0 (73) and EnrichedHeatmap v1.24.0 (82) packages. Box plots and associated Wilcoxon tests were computed using means of bins per gene and were produced using the ggboxplot and stat_compare_means functions, respectively, with the default settings of the ggpubr v0.5.0 package (https://CRAN.R-project.org/package=ggpubr). Publicly available Htz1 ChIP-seq data in wild-type yeast were obtained from Gene Expression Omnibus (GEO) under accession code GSE54105 (48). Following alignment and peak calling as previously described, the intersection of Htz1 genomic regions with Sfp1 peaks was completed using Intervene v0.6.5 (83).

### Carbon shift and glucose pulse assay

Yeast cells were grown in YPD overnight at 30 °C to the exponential phase. The next day, cells were collected by centrifugation and washed twice with water before resuspending in the media containing 2% raffinose as the single carbon source. Glucose was then added to the culture to a final 2% after raffinose incubation. Total RNA was extracted at 15min, 30min and 60min upon glucose addition.

## Data Availability

NGS assays reported in this study are available at the GEO repository under the following accession numbers: ChIP-seq data (GSE267779, Token: kpylmiiibnubbep), RNA-seq data (GSE268202,Token: kxqzaaygzbyrjez), CUT&RUN-seq data (GSE267939, Token: cjizweiyrhgfbgb).

## SUPPLEMENTAL FIGURE LEGENDS

**Figure S1.**
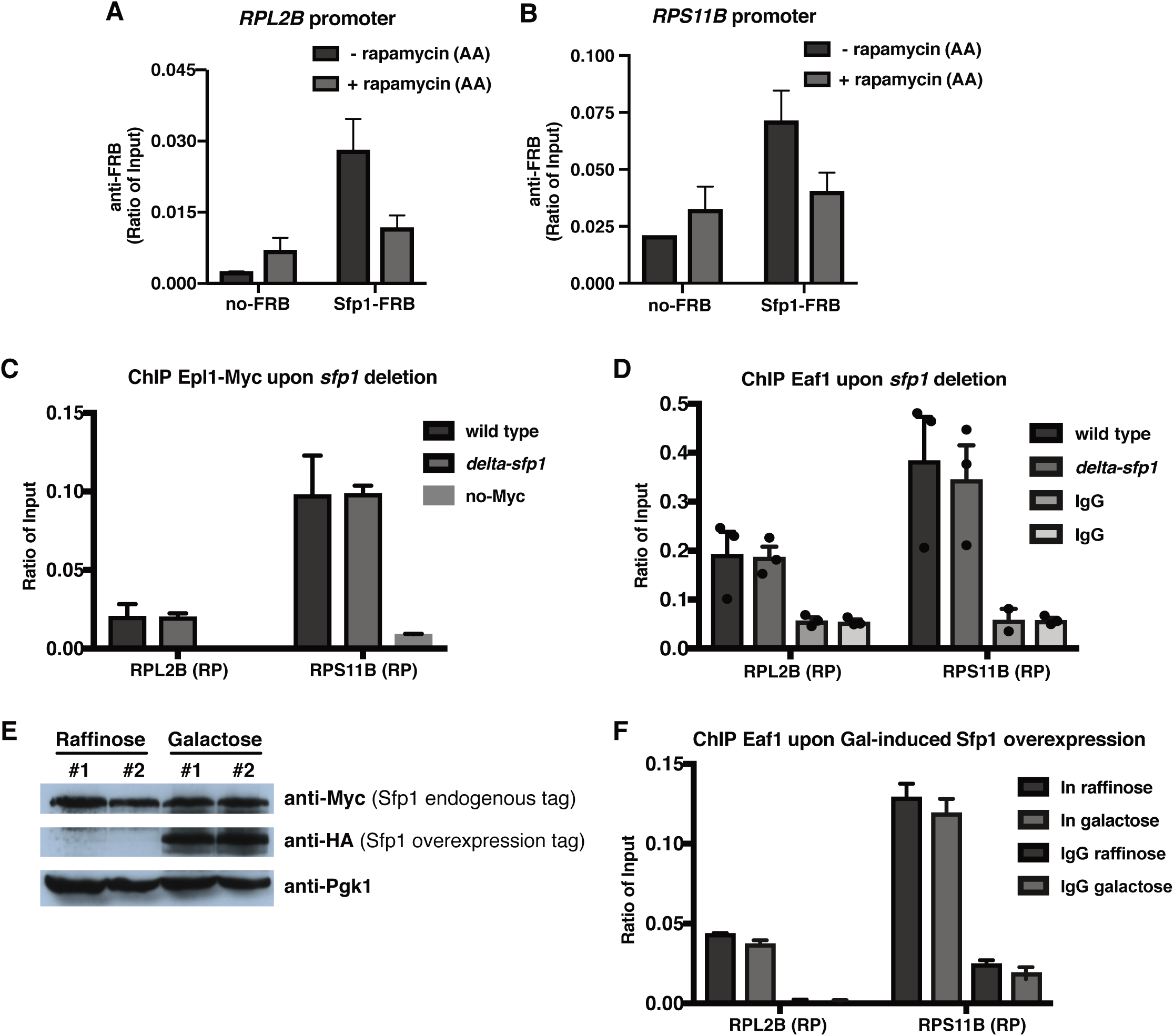
NuA4 binding at RP gene promoters is not affected by Sfp1 depletion. **A-B)** Loss of Sfp1-FRB at *RPL2B* and *RPS11B* gene promoters upon anchor-away treatment determined by ChIP-qPCR using an anti-FRB antibody. Sfp1-FRB or untagged (no-FRB) cells were treated with rapamycin (1 µg/mL) for 2 hours. Error bars represent the range of two independent experiments. **C-D)** ChIP-qPCR showing the binding of the NuA4 complex (Epl1 and Eaf1 subunits) at the promoters of *RPL2B* and *RPS11B*. A *SFP1*-deleted strain shows no significant change in NuA4 binding compared to the wild-type strain. Values are a ratio of input (%). Error bars in D are the standard error of the values (*n*=3). **E)** The Gal-induced Sfp1 overexpression plasmid (HA-tagged Sfp1) was constructed and transformed into the strain expressing endogenous myc-tagged Sfp1. Sfp1 protein levels were determined by western blotting before and after overexpression. Endogenous Myc-tagged Sfp1 shows no protein level change in raffinose and galactose. The Gal-induced Sfp1 overexpression was detected using the anti-HA antibody. **F)** ChIP-qPCR signal showing Eaf1 binding at RP genes (*RPL2B* and *RPS11B*) promoters is not affected upon Sfp1 overexpression. Values are ratio of input and error bars are the range of two independent experiments.

**Figure S2.**
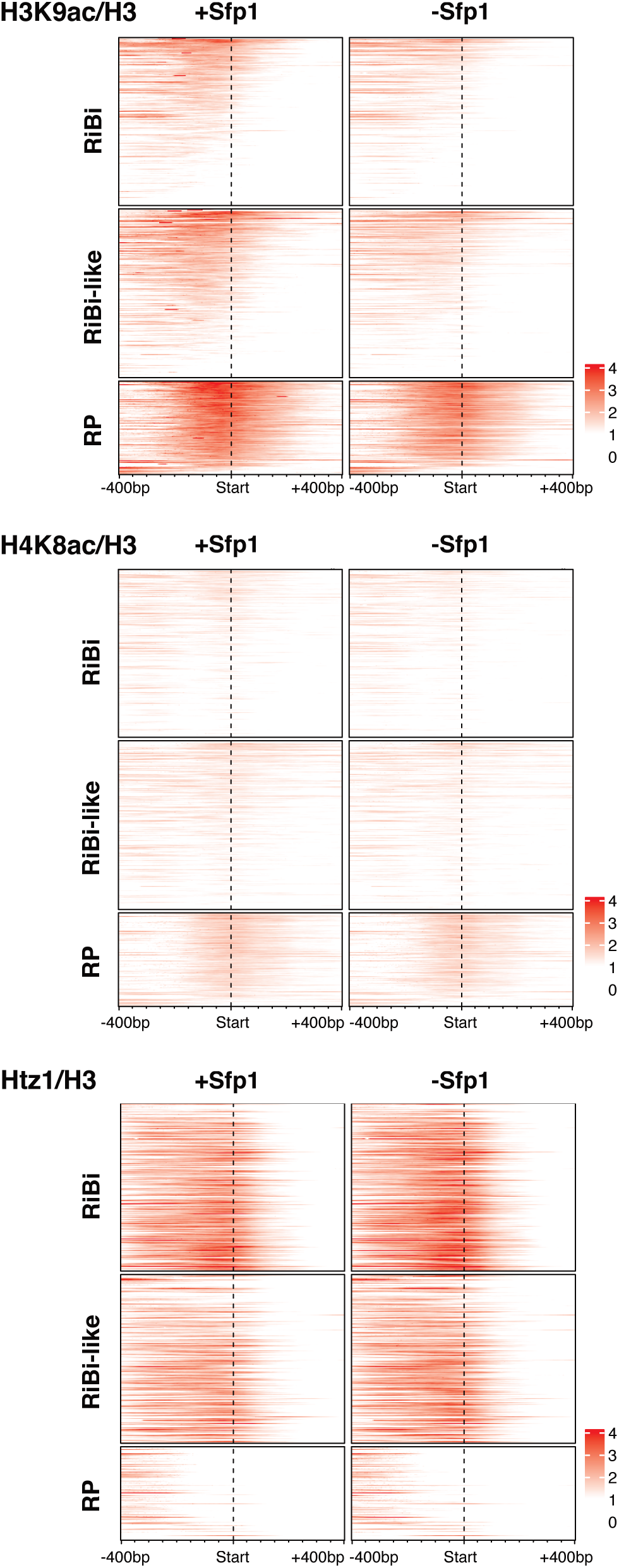
Heatmaps showing histone acetylation and Htz1 occupancy changes upon Sfp1 depletion. ChIP-seq of H3K9ac, H4K8ac and Htz1 occupancy in normal and Sfp1-depleted conditions. An anti-H3 antibody was used to determine total histone occupancy as a control. Signals for a window of −400 to +400 bp relative to TSS are displayed.

**Figure S3.**
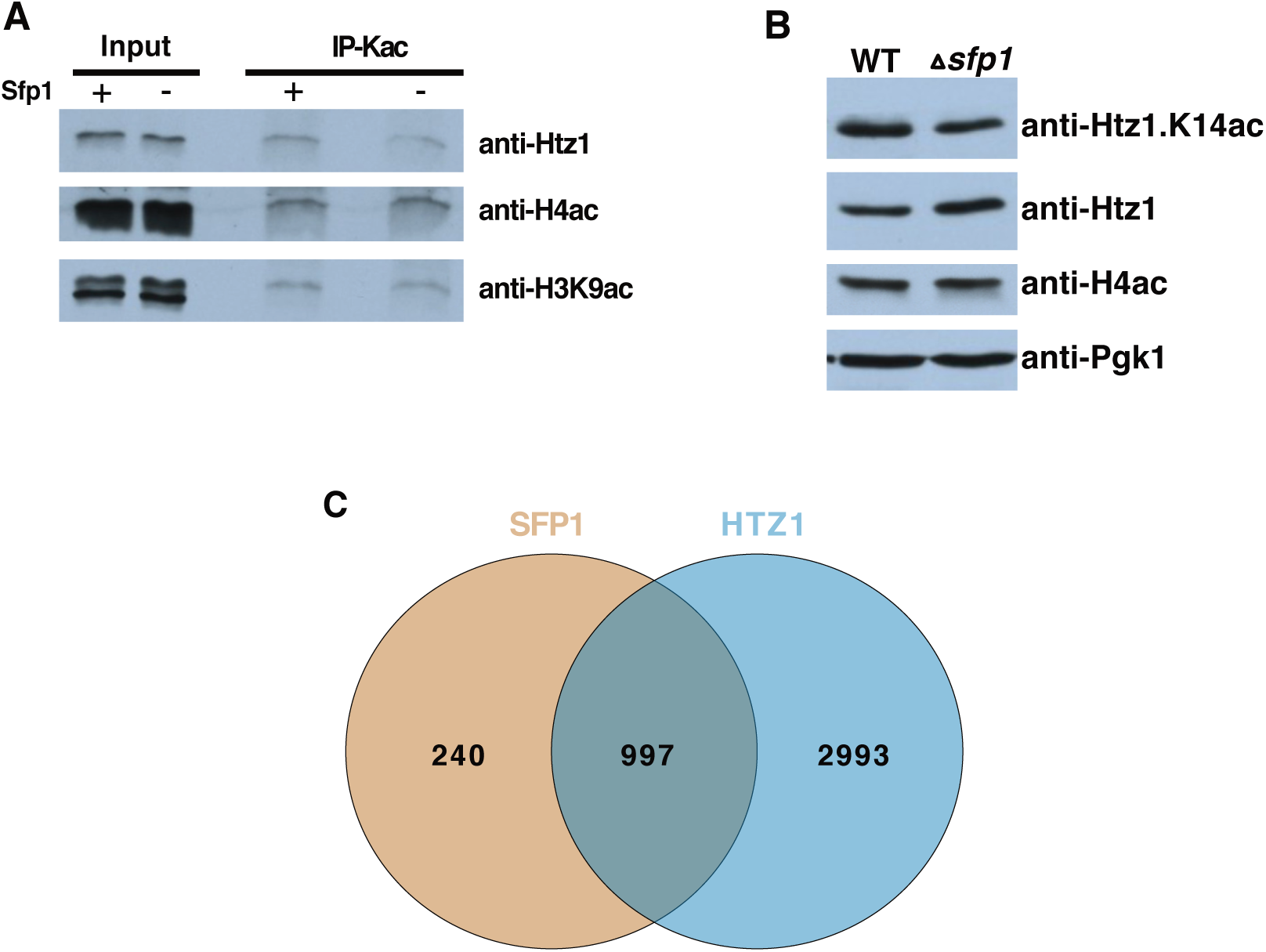
Sfp1 affects histone variant Htz1 occupancy and its acetylation. **A)** Htz1 acetylation is decreased in Sfp1-depleted cells. Immunoprecipitations (IP) were performed with whole cell extracts from cells depleted or not for Sfp1 using an anti-acetyl-lysine antibody. Htz1, H4 and H3 acetylation levels were determined by western blotting. **B**) Western blot analysis of whole cell extracts from wild-type and *sfp1* deleted cells. Bulk histone H4 acetylation, Htz1 and Htz1 acetylation levels are shown. Pgk1 signal is shown as loading control. **C)** Venn diagram showing the overlap between our wild-type Sfp1 CUT&RUN peaks (Fig 3) and published Htz1 ChIP-seq peaks.

**Figure S4.**
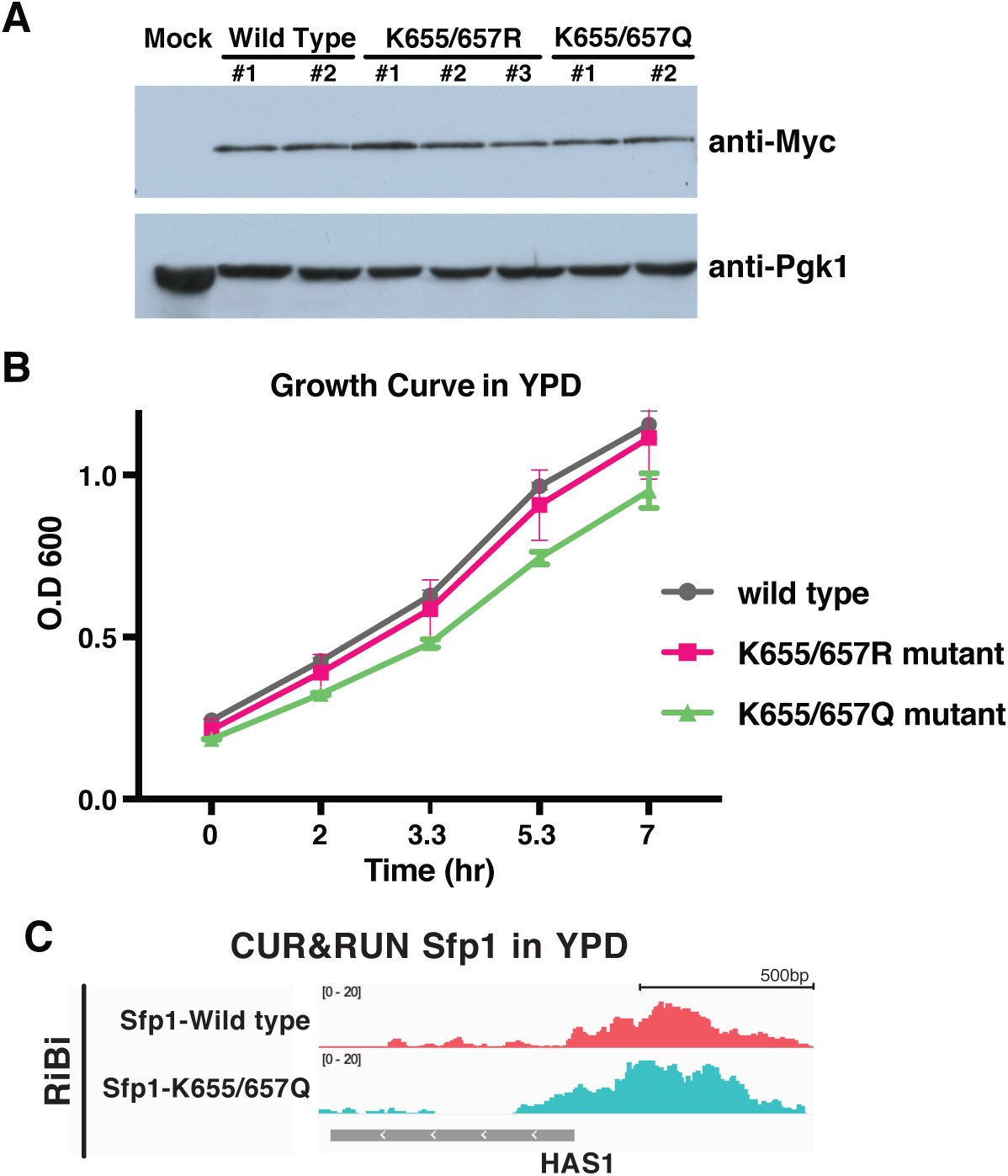
K655/657 mutants show almost no effect on Sfp1 protein expression and slight growth defect in YPD. **A)** Expression of Sfp1 proteins in wild-type and mutated strains. Pgk1 is a reference protein. **B)** The indicated strains were grown to the log phase in YPD. Growth curves are depicted at representative time points. Error bars are standard deviation from three different cultures. **C)** IGV tracks showing Sfp1 CUT&RUN peaks at the selected *HAS1* gene (RiBi) promoter.

**Figure S5.**
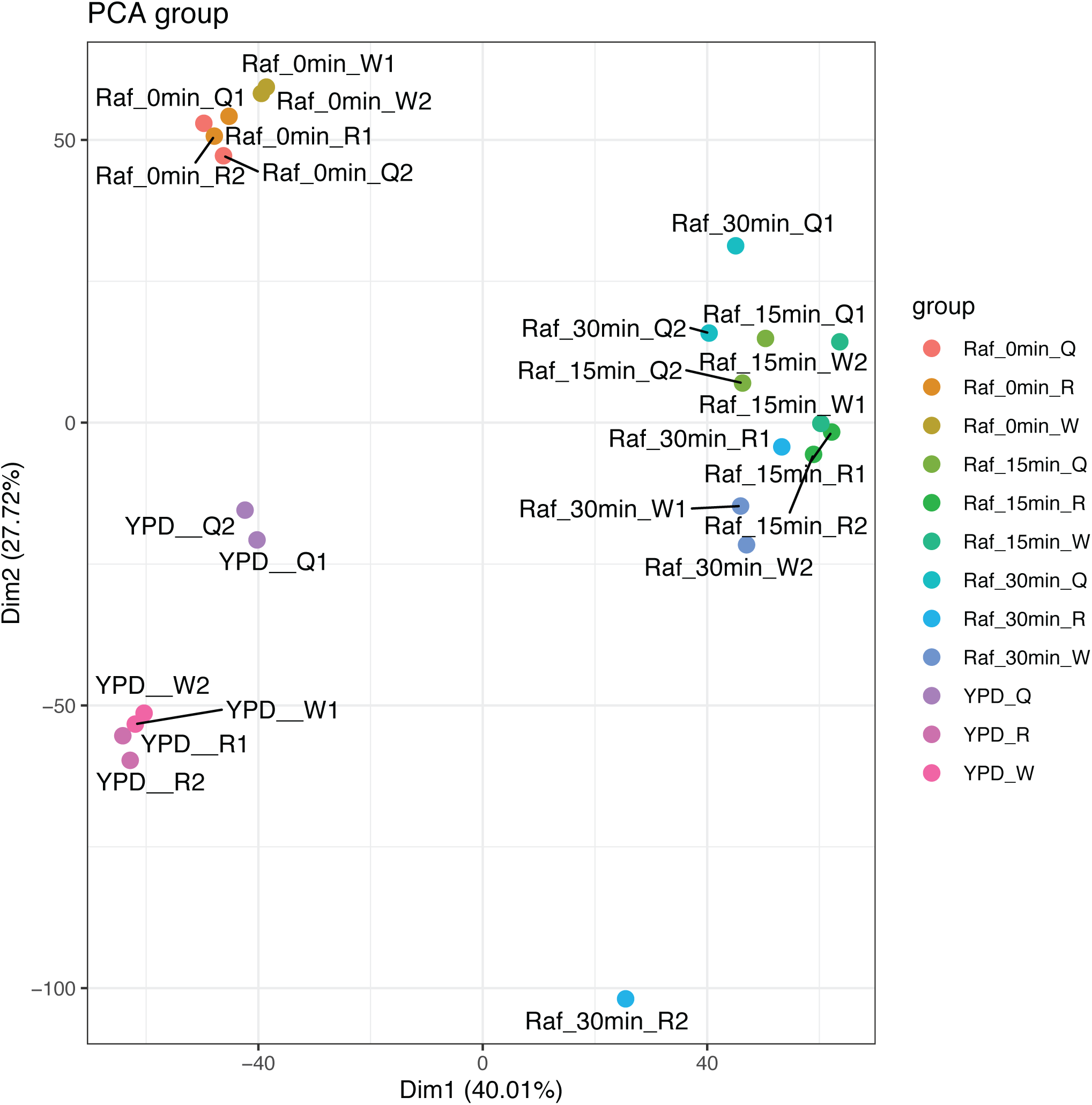
Principal component analysis (PCA) of the RNA-Seq samples. The PCA figure demonstrates variance between RNA-seq samples. Wild type and mutated samples (duplicates) at different time points are represented by different colours, as indicated.

**Figure S6.**
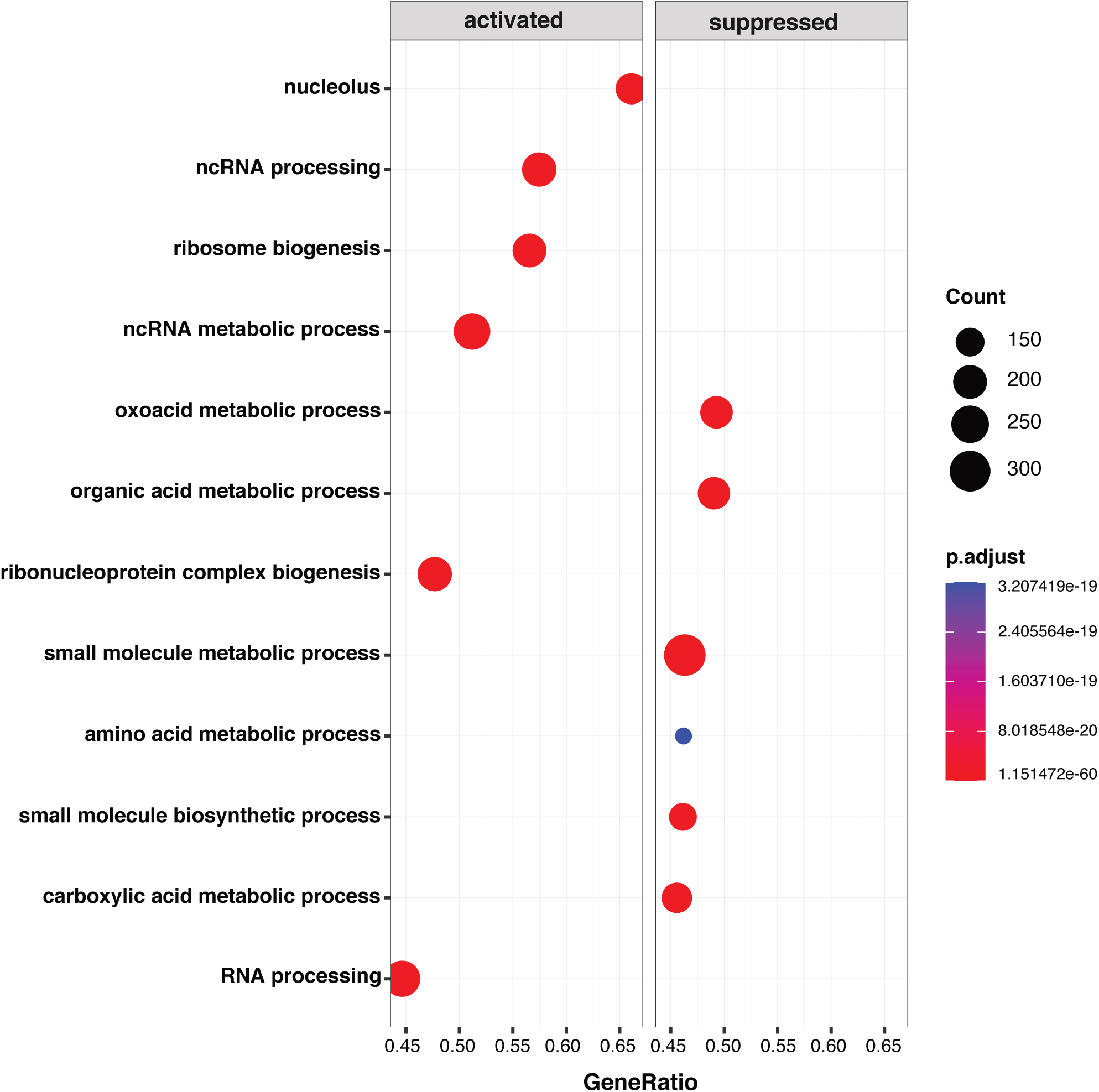
Genes involved in metabolic processes are downregulated in the Sfp1 K655/657Q strain. Gene set enrichment analysis (GSEA) of RNA-seq data was performed on wild-type cells and acetyl-mimic Sfp1 cells growing in YPD. Top up- and down-regulated pathways are shown.

**Figure S7.**
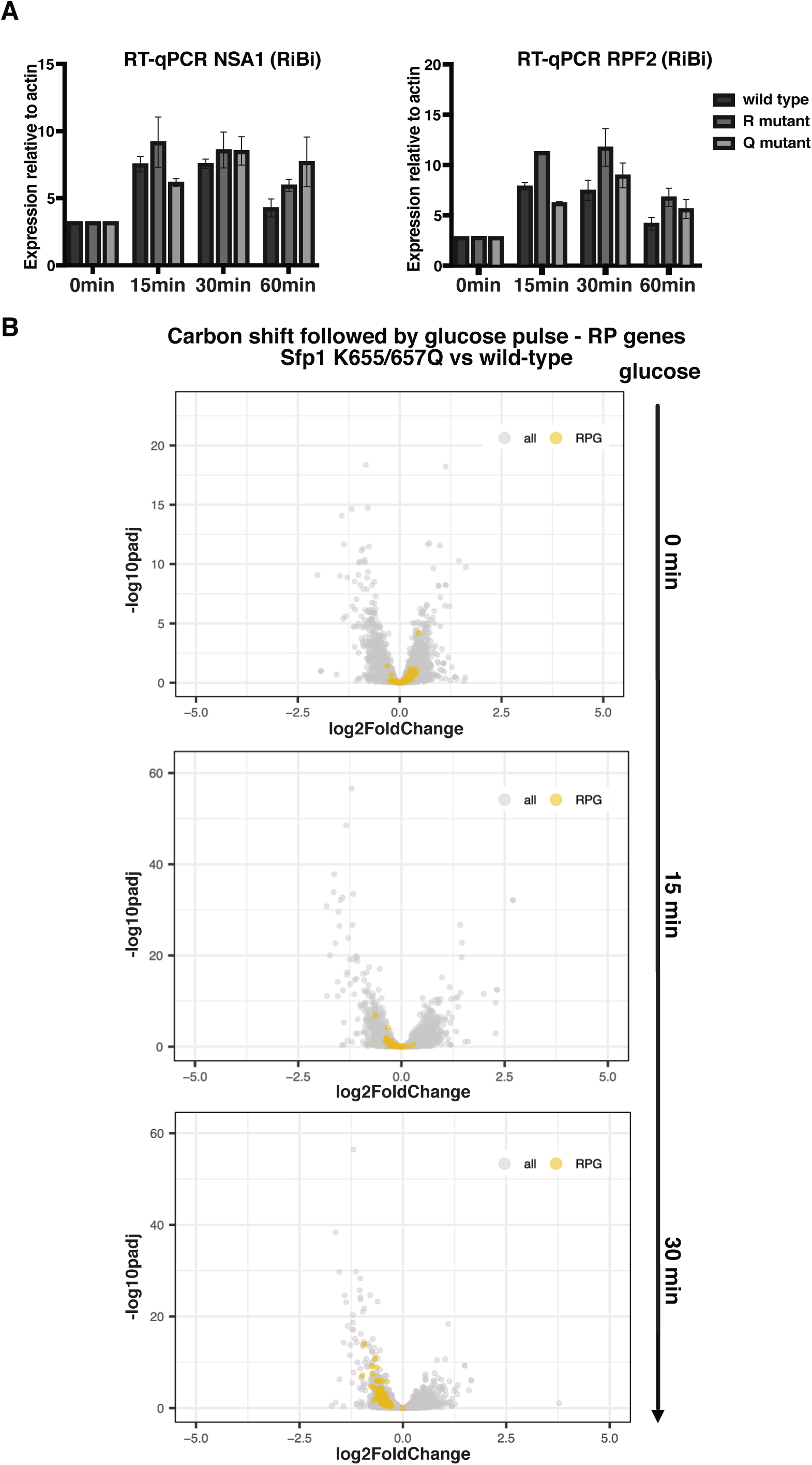
Sfp1 regulates RiBi and RP gene expression through different mechanisms upon glucose pulse after the shift in carbon source. **A)** RNAs were extracted from log phase wild-type Sfp1, K655/657R, and K655/657Q strains grown in indicated times of glucose pulse. Gene expression levels of selected RiBi genes (*NSA1* and *RPF2*) were quantified, relative to levels of *ACT1* mRNA, by RT-qPCR. Values were normalized to time 0 in each strain. Error bars represent the range of two independent experiments. **B)** Volcano plots showing differential gene expression analysis by RNA-seq of cells expressing wild-type Sfp1 and K655/657Q mutation. Expression changes at different time points upon glucose pulse after carbon source shift are shown. RP genes are highlighted.

**Supplemental Table 1.** Yeast strains used in this study.

| Strain | Genotype | Reference |
| --- | --- | --- |
| LPY3431 | *MATa his3Δ200 leu2-3,112 trp1Δ1 ura3-52* | (Clarke et al., Mol Cell Biol 1999) |
| W303-1A | *MATa ade2-1 his3-11,15 leu2-3,112 can1-100 trp1-1 ura3-1* | (Thomas and Rothstein, Cell 1989) |
| JKM139 | *MATa hoΔ hmlΔ::ADE1 hmrΔ::ADE1 ade1-100 leu2-3,112 trp1::hisG' lys5 ura3-52 ade3::GAL::HO* | (Lee et al., Cell 1998) |
| QY3530 | JKM139 *fpr1Δ::HphMX tor1-1::URA3 RPL13A-2\*FKBP12::TRP1* (No-FRB) | (84) |
| QY3572 | Isogenic to QY3530 except *FRB-ESA1::KanMX* | (84) |
| QY3810 | Isogenic to QY3530 except *SFP1-FRB::KanMX* | this study |
| QY3811 | Isogenic to QY3530 except *SFP1-FRB::KanMX EPL1-FLAG::NatRMX* | This study |
| QY3801 | LPY3431 *sfp1Δ::KanMX* | this study |
| QY3802 | LPY3431 *SFP1-MYC::KanMX* | this study |
| QY3803 | LPY3431 *SFP1-MYC::KanMX EPL1-FLAG::URA* | this study |
| QY3804 | LPY *EPL1-FLAG::URA* | this study |
| QY3805 | LPY3431 *SFP1-MYC::KanMX* + pBG1805(GAL-Sfp1-6*HIS-HA) | this study |
| QY3806 | LPY3431 *SFP1-K655R-K657R-MYC::KanMX* | this study |
| QY3807 | LPY3431 *SFP1-K655Q-K657Q-MYC::KanMX* | this study |
| QY3808 | LPY3431 *SFP1-K655R-K657R* | this study |
| QY3809 | LPY3431 *SFP1-K655Q-K657Q* | this study |
| YMD070 | *MATα leu2Δ0 ura3Δ0 his3-1 met15Δ0 sir2Δ::HYGMX hst1Δ::KANMX hst2Δ::HIS3MX* | (84) |
| QY3812 | Isogenic to YMD070 except *SFP1-MYC::KanMX* | this study |
| QY3813 | Isogenic to YMD070 except *SFP1- K655R-K657R-MYC::KanMX* | this study |
| QY3814 | Isogenic to YMD070 except *SFP1- K655Q-K657Q-MYC::KanMX* | this study |

